# Complementary vertebrate *Wac* models exhibit phenotypes relevant to DeSanto-Shinawi Syndrome

**DOI:** 10.1101/2024.05.26.595966

**Authors:** Kang-Han Lee, April M Stafford, Maria Pacheco-Vergara, Karol Cichewicz, Cesar P Canales, Nicolas Seban, Melissa Corea, Darlene Rahbarian, Kelly E. Bonekamp, Grant R. Gillie, Dariangelly Pacheco-Cruz, Alyssa M Gill, Hye-Eun Hwang, Yeong-Eun Kim, Katie L Uhl, Tara E Jager, Marwan Shinawi, Xiaopeng Li, Andre Obenaus, Shane Crandall, Juhee Jeong, Alex Nord, Cheol-Hee Kim, Daniel Vogt

## Abstract

Monogenic syndromes are associated with neurodevelopmental changes that result in cognitive impairments and neurobehavioral phenotypes including autism and seizures. Limited studies and resources are available to make meaningful headway into the underlying molecular mechanisms that result in these symptoms. One such example is DeSanto-Shinawi Syndrome (DESSH), a rare disorder caused by pathogenic variants in the *WAC* gene. Individuals with DESSH syndrome exhibit a recognizable craniofacial gestalt, developmental delay/intellectual disability, neurobehavioral symptoms that include autism, ADHD, behavioral difficulties and seizures. However, no thorough studies from a vertebrate model exist to understand how these changes occur. To overcome this, we developed both murine and zebrafish *Wac/wac* deletion mutants and studied whether their phenotypes recapitulate those described in individuals with DESSH syndrome. We first show that the two *Wac* models exhibit craniofacial and behavioral changes, reminiscent of abnormalities found in DESSH syndrome. In addition, each model revealed impacts to GABAergic neurons and further studies showed that the mouse mutants are susceptible to seizures, changes in brain volumes that are different between sexes and relevant behaviors. Finally, we uncovered transcriptional impacts of *Wac* loss of function in mice that will pave the way for future molecular studies into DESSH. These studies present two new animals that begin to uncover some biological underpinnings of DESSH syndrome and elucidate the biology of *Wac*.

## Introduction

Pathogenic variants in the WW domain-containing adaptor with coiled coil, *Wac*, gene cause a neurodevelopmental disorder called DeSanto-Shinawi (DESSH) syndrome (aka WAC-related disorder) [1]. DESSH syndrome is characterized by a constellation of developmental delay/intellectual disability, cranio-facial dysmorphism, hypotonia, seizure, gastrointestinal problems (e.g., constipation and gastroesophageal reflux), and ophthalmological abnormalities (e.g., refractive errors and strabismus) [1]. In addition, individuals with DESSH syndrome exhibit arrays of neurobehavioral phenotypes including aggression, attention deficit hyperactivity disorder (ADHD), autism spectrum disorder (ASD) and anxiety [1–5]. *Wac* is a known autism risk gene due to the enrichment of pathogenic genetic variants in ASD cohorts [6]. However, little is known about the function of WAC in the brain and whether suitable vertebrate models could explore the underlying biology of DESSH syndrome phenotypes.

The *Wac* gene encodes a protein with WW and coiled-coil domains, common protein/protein interaction regions. A recent study assessed the role of evolutionarily conserved protein domains in the human WAC protein that uncovered an amino terminal nuclear localization signal as well as a highly conserved disorganized region that contains several putative phosphorylation motifs that are mutated in humans with known *WAC* variants [7]. While the functional role of these domains and of WAC itself are still poorly understood in vertebrates, a multitude of roles for the WAC protein have been proposed, including positive regulation of mammalian target of rapamycin (mTOR) signaling, mitosis, transcription and autophagy [8–11], which were worked out in *drosophila* and immortalized cell lines. However, whether these roles are conserved in vertebrates and their relevance to human DESSH symptoms is still unknown.

To better understand *Wac*’s role in vertebrates and in particular brain-relevant changes, we generated two species-specific vertebrate deletion models to assess whether *Wac* depletion could recapitulate symptoms associated with DESSH syndrome. The rationale for choosing mouse and zebrafish models were: 1) Both are vertebrates; 2) Unique genetic approaches in each model; 3) Species-specific phenotypic screening advantages; 4) Finally, utilizing two distinct vertebrate models to study key brain phenotypes will be a powerful means to uncover the fundamental biology of DESSH syndrome in a manner that is likely to reveal novel neurological underpinnings.

Overall, our models revealed some similarities in core DESSH symptoms, including craniofacial dysmorphism and relevant behaviors implicating decreased learning and social interaction. Notably, while each model displayed common decreased GABAergic markers, there were slight differences between species in targets. Additionally, mice had increased seizure susceptibility, while zebrafish did not; mice had only minor changes to sociability while zebrafish showed a strong change. We further took advantage of the mouse model, which more closely resembles humans, to probe into potential unexplored phenotypes and molecular changes. Despite some differences, both models revealed that males were more susceptible in multiple phenotypes and that male mice exhibited novel molecular changes that could be further investigated. The first vertebrate disease models for DESSH syndrome are presented here, and the multiple shared and unique phenotypes that could begin to inform the underlying biological mechanisms of DESSH.

## Results

### Validation of murine *Wac* and zebrafish *wac* mutants

We obtained sperm from the International Mouse Phenotyping Consortium (IMPC); the *Wac* murine locus on chromosome 18 was genetically modified to include flanking *loxP* (Flox) sites of exon 5 (Figure 1A). Sanger-sequencing of the genotyped PCR validated the locus and an introduced frameshift that resulted in a premature stop codon (Figure 1B). *Wac flox* founders were bred and crossed to *beta-actin-Cre* mice, which express *Cre* in germ cells [12]. We generated wild type (WT) and constitutive heterozygous (Het) progeny of similar size (Figure 1C) but not any live homozygous knockouts. Findings were consistent with data from the IMPC; constitutive *Wac* KOs are embryonic lethal. Results of genotyping using PCR are shown in Figure 1D. WAC protein was also reduced by 64% in Het brains, (shown later in Figure 5).

**Figure 1:**
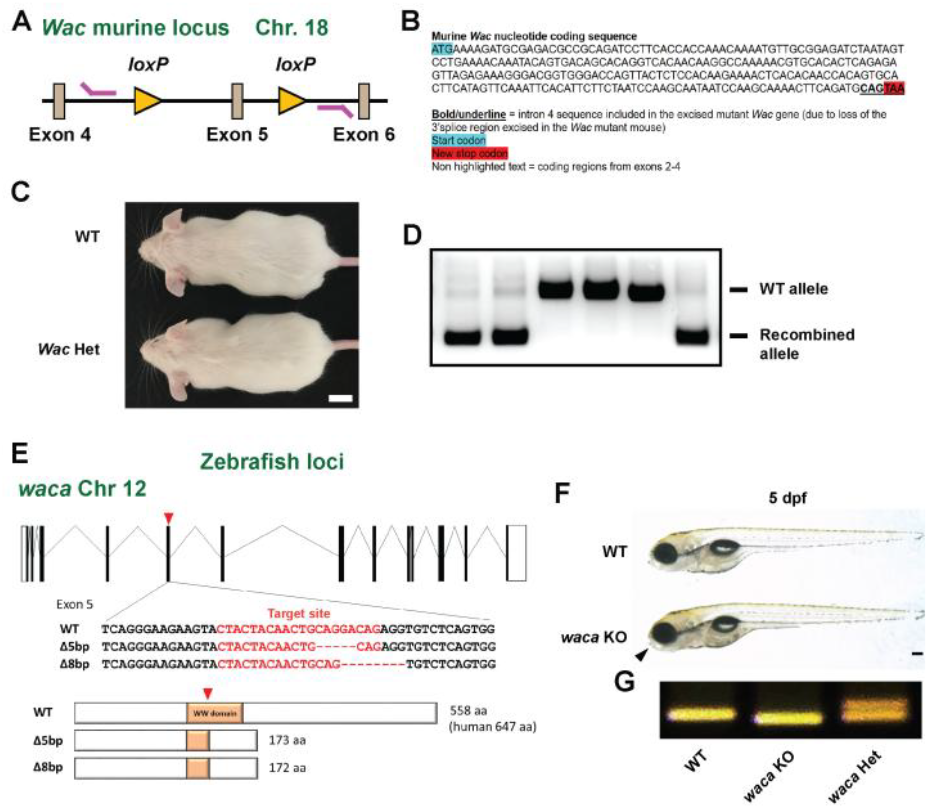
Generation and validation of murine *Wac* and zebrafish *waca* mutants. (A) Floxed mouse *Wac* genetic locus and genotyping primers. Genotyping primers (magenta lines) reside external to the loxP sites (orange triangles) flanking exon 5 and nearby introns. (B) The recombined DNA band was Sanger-sequenced and results show the start codon (blue) and the novel stop codon introduced by the frameshift mutation resulting from Cre-mediated deletion of the locus used to generate the locus. (C) Example images of adult WT and *Wac* Het mice. (D) DNA gel of genotyping for WT and recombined (Het) *Wac* alleles. (E) *waca* KO zebrafish were generated by CRISPR/Cas9. KO target sites for sgRNAs are indicated by red arrowheads and predicted protein structures for KO mutations indicated below panel. (F) WT and *waca* KO zebrafish have grossly normal sizes but *waca* KO exhibited a shortened jaw (black triangle) at 5 dpf. (G) example *waca* genotyping results. Abbreviations: (aa) amino acid; (bp) base pair; (Chr) chromosome; (dpf) days post fertilization; (WT) wild type. Scale bars (C) = 1 cm and (F) = 200 µm.

Zebrafish harbor two *wac* genes on chromosome 12, *waca*, and 2, *wacb*. A CRISPR/CAS9 approach was used to target these loci to create *waca* and *wacb* KOs and then interbreed to generate *waca*; *wacb* double KOs. Notably, *waca* expression was more widespread and higher in the brain of zebrafish compared to *wacb* (Figure S1A), indicating that *waca* might be more important for DESSH. The *waca* mutant had a deletion in coding exon five (Figure 1E) with *waca* progeny grossly normal in size by 5 days post fertilization (dpf) but with a shortened lower jaw (Figure 1F); genotyping results are shown in Figure 1G. The *wacb* KO and *waca*;*wacb* double KO zebrafish are shown in Figure S1B. Since the *wacb* KOs did not show phenotypes like the *waca* mutants, we did not report further data. Moreover, since double KOs exhibited early lethality we chose to solely focus on *waca* KOs for this report. Some possible causes of lethality are denoted in Figure S1B, with double KOs showing evidence of heart edema and loss of the swim bladder.

### Craniofacial changes in murine and zebrafish mutants

Craniofacial dysmorphism is a cardinal feature in patients with DESSH syndrome [1–3, 5, 13–17], with larger frontal facial features being common, including larger foreheads. We first wondered whether a reduction in *Wac* dosage in mice could recapitulate this finding. Postnatal day (P)0 and P30 mouse skulls were examined. During neonatal ages, a widening of the fontanels and sutures along the midline of the calvaria was noted (Figure 2A, B). While both the anterior and posterior regions of the calvaria were affected, the anterior region had the greater change (Figure 2A, B, E, F with pseudo-colored areas in 2A’, B’; anterior p = 0.0001, posterior p = 0.01). By P30, there remained a noticeable gap in the interfrontal suture of the *Wac* Het. Concomitantly, the width of the skull across the frontal bones was significantly increased in *Wac* Hets, while the width across the parietal bones was normal (Figure 2C, D, G, H, p = 0.002 Frontal width). A feature that was notable was a shortened lower jaw in the *waca* KO zebrafish compared to WTs at 10 dpf (Figure 2I-K, p = 0.006), suggesting similar abnormalities in frontal craniofacial structures. In addition, the *waca* mutants also exhibited changes in other frontal craniofacial features, including an increased width spanning Meckel’s cartilage at 13 dpf (Figure 2L-N, p = 0.02). The *waca* KO zebrafish survived into adulthood and continued to show morphological craniofacial defects of shortened lower jaw, sunken head and broad nasal tip at adult stage (data not shown). Thus, both models exhibit craniofacial changes relevant to DESSH syndrome.

**Figure 2:**
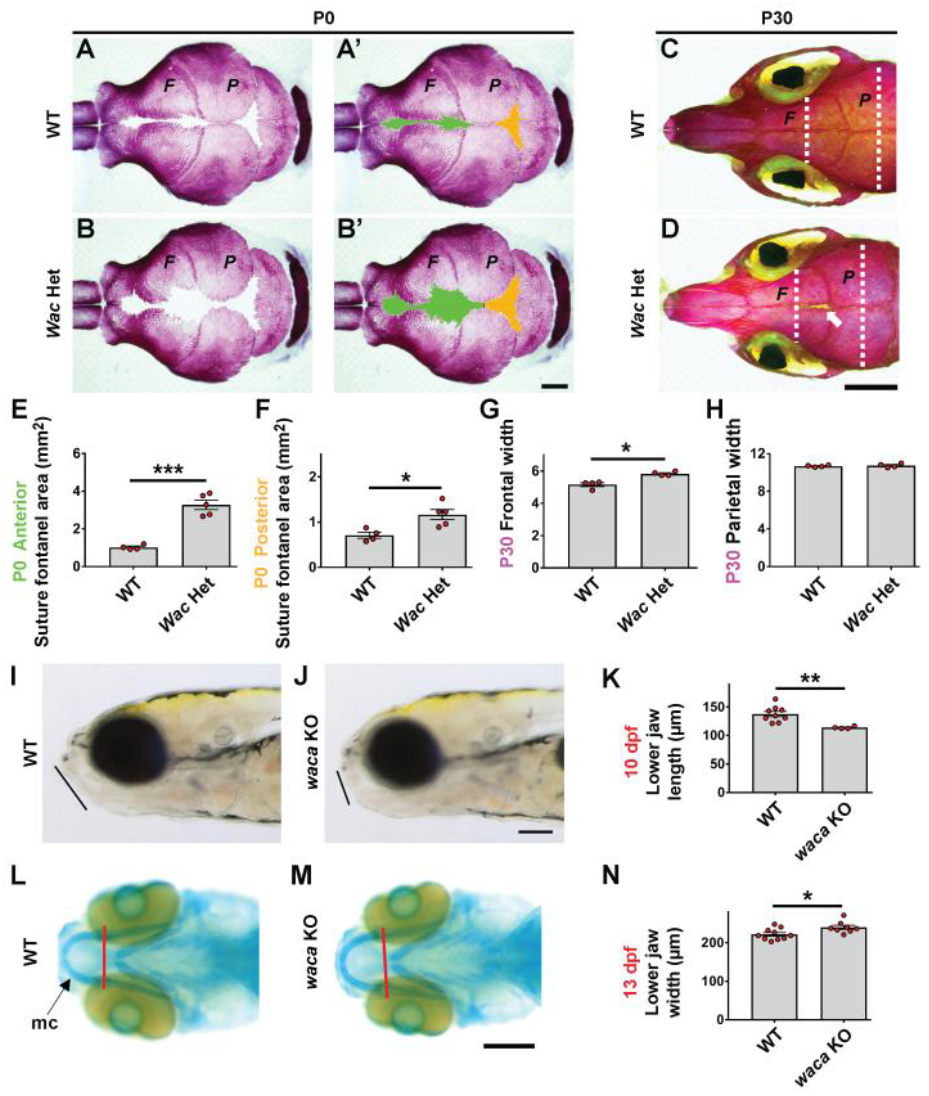
Craniofacial changes in mutants. (A, B) Dorsal views of the P0 calvaria stained with Alizarin red for bone. (A’, B’) The suture and fontanel areas in (A) and (B) are pseudo-colored green (anterior) and orange (posterior). (C, D) Dorsal views of P30 WT and *Wac* Het skull bones; arrow (D) points to a gap in the interfrontal suture. (F) Frontal bone and (P) Parietal bone; white dashed lines are widths measured in (G, H). (E, F) Quantification of P0 fontanel suture areas pseudo colored in A’ and B’; WT n = 4, *Het* n = 5. Quantification of the skull width across the frontal (G) and parietal (H) bones at P30 in WTs and *Wac* Hets; WT n = 4, Het n = 4. (I, J) The *waca* KO zebrafish show a shortened jaw structure, compared to WT. Lines denote lower jaw length. (K) Quantification of lower jaw length; n = 9 WTand 4 *waca* KO. (L, M) Cartilage staining of zebrafish using alcian blue, ventral view. (N) Measurements of the width of Meckel’s cartilage in *waca* KO zebrafish at 13 dpf (red line); WT n = 10 and KO n = 8. Data are expressed as the mean ± SEM. *p < 0.05, **p < 0.01 and ***p < 0.001. Scale bars (B’) = 1 mm, (D) = 4 mm, (J, M) = 200 µm.

### Behavioral changes in *Wac/wac* mutants

Loss of *Wac* may impact social/cognitive behavior, as seen among individuals diagnosed with DESSH syndrome and as previous models exhibit intellectual challenges [3, 5, 14, 15, 17]. We noticed that Het mutant mice had normal ambulation, total distance traveled and time spent in different areas of the open field (Figure S2A). In a 3-chamber social test, while some *Wac* Hets trended towards less social discrimination using a previously described test [18], they did not significantly differ from WTs (Figure 3A). However, *Wac* Hets did show significant impairments in the Y-maze that tests working memory [19, 20] (Figure 3B, p = 0.047). Anxiety (elevated plus maze) and spatial memory (radial arm water maze) were not grossly changed but similar to the social discrimination test, cohorts of *Wac* Hets trended towards impairment in each test (Figure S2B, C).

**Figure 3:**
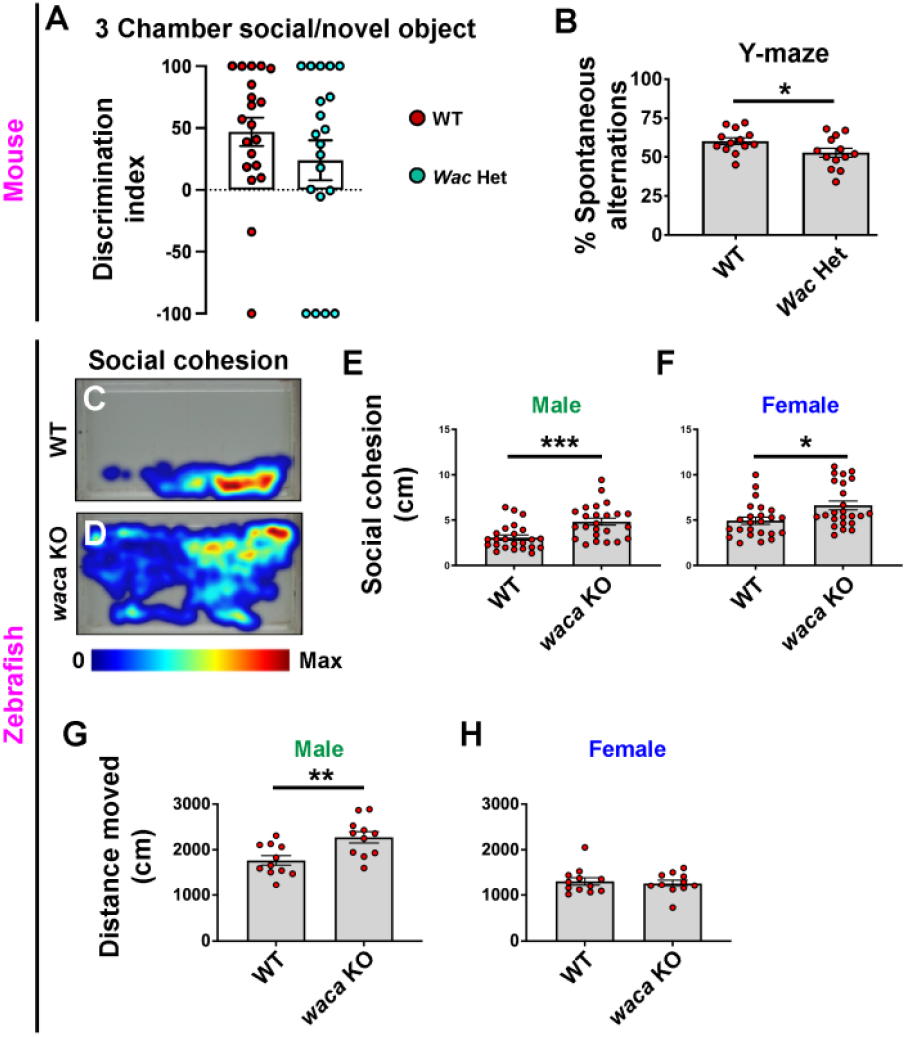
Behavioral changes in mutants. (A) 6-8 week WT and *Wac* Het mice were tested in the 3-chamber social/novel object test, WT n = 20 and Het n = 20, and the Y-maze (B), n = 13 both groups, with *Wac* Hets showing deficits in the Y-maze. (C, D) Example heat maps during the 17-19-minute timeframe of the social cohesion test in zebrafish; *waca* KO males and females were more dispersed (E, F), see methods for test details. (G, H) Distance moved was greater in male *waca* KOs in the novel tank assay, n = 11, both groups, compared to females, n = 12 (WT) and 11 (KOs). Data are expressed as the mean ± SEM; *p < 0.05, **p < 0.01 and ***p < 0.001.

We next tested zebrafish in a social cohesion assay; WT fish stayed in close contact while *waca* KOs were dispersed during the trial, see heatmap distribution of zebrafish in Figure 3C, D. While both male and female zebrafish *waca* KOs showed deficits in this assay, we report the sexes independently since WT males and females had significant differences in baseline social cohesion. Moreover, male zebrafish *waca* KOs had a greater change in social cohesion than females (Figure 3E, F, p = 0.0007 males, p = 0.01 females). In a novel tank assay to test hyperactivity, male *waca* KO fish moved greater distances, throughout the test compared to WTs (Figure 3G, p = 0.005), while there was no significant differences in females (Figure 3H). While differences exist in zebrafish locomotion/hyperactivity, both models share deficits in some behaviors.

### Elevated seizure susceptibility and depleted GABAergic markers in *Wac* heterozygous mice

The previous behavioral changes suggested that core neuron cell ratios might be altered, potentially through an imbalance in GABAergic/glutamatergic neurons that could impact this and other disorders [21, 22]. Epilepsy occurs frequently in individuals with DESSH syndrome [1, 3, 4, 15, 17], thus, we first tested if loss of *Wac* expression was sufficient to increase seizures in our mouse model. We assessed whether the GABAA receptor antagonist, pentylenetetrazol (PTZ), could induce seizures in mice with *Wac* loss of function. The drug was administered intraperitoneal at a subthreshold dose (50mg PTZ per kilogram of mouse body weight) that rarely elicited a tonic-clonic seizure in WTs. We hypothesized that if loss of *Wac* promoted a shift favoring brain excitation/inhibition then *Wac* hets would exhibit more severe seizure behaviors by P30.

PTZ was administered and mice were observed for 20 minutes (Figure 4A); most mice fully recovered by 15 minutes and all WT mice did not exhibit any behavioral changes by twenty minutes post injection. Behaviors included freezing, motor twitches, loss of balance, seizures and hyperactivity, which are detailed in the methods. While PTZ led to subtle behavioral changes in WT mice but rarely elicited a seizure. However, *Wac* Hets exhibited elevated seizure behaviors, including hunched body posture, arching tail and forelimb clonus as well as succumbing to seizures (Figure 4B, p < 0.0001). Thus, loss of *Wac* leads to increased seizure susceptibility in mice, consistent with one of the comorbidities observed in ∼40% of individuals with DESSH syndrome. In contrast, when zebrafish were tested for seizure susceptibility, the *waca* KOs did not show any gross changes although both WTs and mutants responded to PTZ (data not shown), suggesting that this specific symptom was not conserved between the 2 vertebrate models.

**Figure 4:**
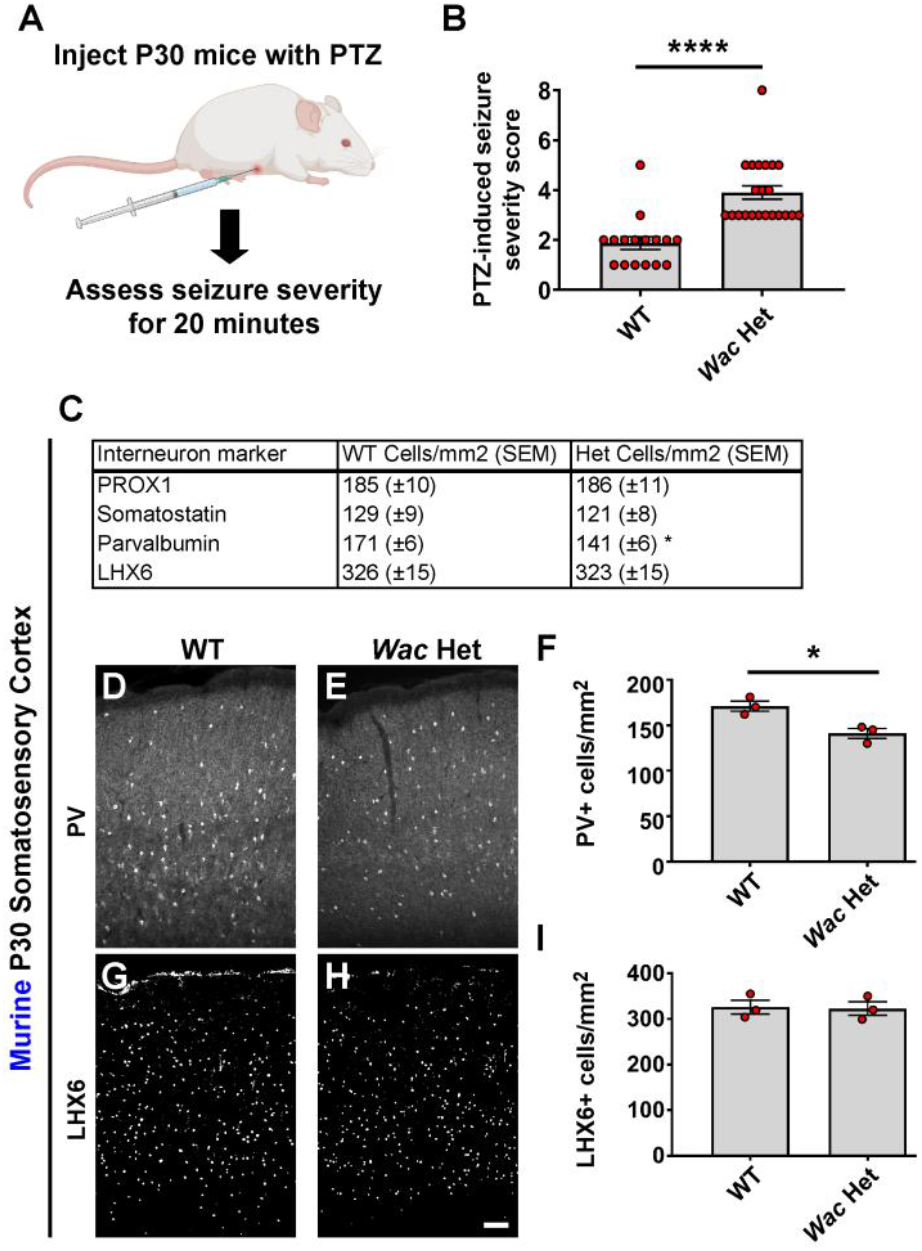
Murine *Wac* depletion leads to elevated seizure susceptibility and loss of GABAergic markers. (A) Schema depicting the seizure induction by PTZ; mouse image was made by BioRender software. Briefly, P30 mice were administered PTZ intraperitoneally and then assessed for the highest seizure severity score over the course of 20 minutes. The maximum seizure severity score over 20 minutes was quantified (B); n = 16 (WTs) and 22 (Hets). (C) Table of cortical interneuron cell counts in the somatosensory cortex at P30. (D-F) Immunofluorescent images of PV and cell density quantification in mice somatosensory cortex, n = 3, both groups. (G-I) Immunofluorescent images of LHX6 and cell density quantification in mice somatosensory cortex, n = 3, both groups. Data are expressed as the mean ± SEM, *p < 0.05 and ****p < 0.0001. Scale bar = 100µm (H).

### Some GABAergic deficits in mouse and zebrafish *Wac/wac* mutants

While we only found a susceptibility to seizures in mice, an imbalance in excitatory and inhibitory cell populations may still exist in each model but with differential impacts on seizures and/or behaviors. One possibility is a decrease in GABAergic cortical interneuron (CIN) function, a diverse group of inhibitory neurons whose dysfunction has been implicated in other syndromes with epilepsy and/or autism [23–28]. To assess whether *Wac* may impact CIN populations, we assessed 3 broad groups of CINs; parvalbumin (PV)+ and somatostatin (SST)+, which label ∼70% of GABAergic interneurons [29], and prospero homeobox 1 (PROX1) expressing, which labels the majority of caudal derived interneurons [30]. The cell density of SST and PROX1 CINs were normal, but there was ∼18% decrease in PV+ CINs in the Het (Figure 4C-F, p = 0.02); example images of SST and PROX1 cells are shown in Figure S5. To probe whether PV expression is due to loss of cells or simply the expression of PV, we assessed for the CIN marker, LHX6, which is expressed in all PV+ CINs in the somatosensory cortex [28]. We found no decrease in LHX6 cell numbers (Figure 4C, 4G-I), suggesting this phenotype is likely due to impacts upon PV expression rather than loss of this group of interneurons.

We also studied GABAergic markers in *waca* KO zebrafish. We first probed via in situ for the various parvalbumin transcripts in zebrafish but found no differences, including *pvalb6*, which is the only one expressed in the brain (Figure S3). We next looked at the pan GABAergic gene, *gad1b*, in the forebrain and compared *gad1b* expression in WTs and *waca* KOs. Notably, we uncovered a significant reduction in forebrain area that expressed the transcript (Figure S4, p<0.0001); WT and *waca* KO zebrafish brains were not altered in general size, suggesting this measurement was not biased due to a smaller or larger brain due to genotype. These and previous data suggest that GABAergic neuron populations are impacted by mouse and zebrafish *Wac*/*waca* loss of function and potentially contribute to DESSH syndrome.

### Normal numbers of major brain cell types and proteins in *Wac* mutant mice brains

Since many antibodies are not available for zebrafish, and mice are closer to humans in gene conservation, we chose to focus on our mouse model to next assess if any other major brain cell populations, signaling events (including post-translational modifications) or gross synaptic markers were altered by loss of *Wac*. Thus, we first performed cell density counts for different classes of neuronal cells in our mouse model that included neurons, oligodendrocytes, astrocytes and microglia in P30 somatosensory cortices (Figure 5A). The total number of neurons was assessed by labeling for NeuN, while oligodendrocytes, astrocytes and microglia were labeled with OLIG2, S100beta and IBA1, respectively; representative images of these markers are shown in Figure S5. No differences in these major cell types were observed between the genotypes in mice.

Next, we assessed mouse P30 somatosensory cortices for proteins changed in other *Wac* models or markers of brain synapses. As mentioned earlier, WAC protein was reduced 64% in the Hets (Figure 5B, C, p = 0.002). Despite previous reports in other models [8, 10], we did not observe differences in mTOR activity or ubiquitination of histone 2B. Key pathways including Wnt/beta-catenin and synaptic markers, PSD95 (excitatory synapses) and Gephyrin/Gad65/67 (inhibitory synapses), were not altered (Figure 5B, C). For the MAPK signaling pathway, there was a decrease in active ERK1, 38% and ERK2, 33%, in the *Wac* Hets (Figure 5B, C, pERK1 p = 0.01, pERK2 p = 0.02), suggesting that while major cell types and proteins involved brain function are not altered, some signaling events may be candidates for future studies to understand WAC function.

### Subtle pyramidal neuron lamination but normal glutamatergic electrophysiological properties in mouse *Wac* Hets

Our data thus far revealed altered GABAergic neuron marker expression but no gross changes in other generic neuron numbers, glia numbers or synaptic protein abundance. However, these observations could still be true while subtypes of excitatory neurons in distinct cortical lamina or their physiological/synaptic properties are yet altered. Thus, we also assessed glutamatergic neuron laminar markers in the somatosensory cortices of P30 mice. SATB2, CTIP2 and TBR1 were labeled and the proportion of expressing neurons assessed in layers 2/3/4, 5 and 6 at P30. While there were no significant laminar changes in the Het for SATB2 and CTIP2, there was a proportional increase of TBR1+ cells in layer 6 (Figure S6A-I, p = 0.047). We also examined the intrinsic membrane properties of layer 2/3 neurons in the primary somatosensory cortex of adult WT and Het mice. Overall, there were no gross differences in passive or active membrane properties between *Wac* Het and WT cells (Figure S6J, K and Table S1), suggesting excitatory cortical neurons in the mutants have normal physiological properties. We also assessed synaptic function by measuring spontaneous excitatory postsynaptic currents (EPSCs) in layer 2/3 neurons. We found no differences between WT and *Wac* Het excitatory neurons (Figure S6L, M), suggesting that local excitatory neurons in the neocortex are not measurably impacted by *Wac* depletion in the Het mice.

### Increased brain volume in *Wac* Het mice with a male bias

While our zebrafish model is similar to the mouse in many core symptoms that presents in humans, the *Wac* Het mice recapitulate almost all DESSH syndromes yet tested. Thus, we wanted to take advantage of this model to understand if there may be more changes to the mouse Het brain that could underlie core and other symptoms relevant to humans. Since craniofacial symptoms exist in nearly all DESSH patients and our models have these same alterations, we hypothesized that alterations to the skull shape may also be correlated with altered brain volume. To determine what regions of the mutant mouse brains may be anatomically different, we performed magnetic resonance imaging (MRI) on P30 fixed WT and *Wac* Het brains of both sexes. We first assessed total brain volume of the WT and Hets and found that male *Wac* mutants exhibit a larger whole brain volume than WT males; there were no significant differences in female brain volumes (Figure 6A, p = 0.04). Next, we examined specific regional volumetric changes, starting with cortical domains. Several alterations were observed in both males and some in females, with most cortical regions showing an increase in volume compared to WTs. Notably, only males had increased retrosplenial cortices, while females showed increases in parietal cortices (Figure 6B, retrosplenial cortex male Het vs male WT p = 0.002 and female WT vs. female Het p = 0.01).

**Figure 5:**
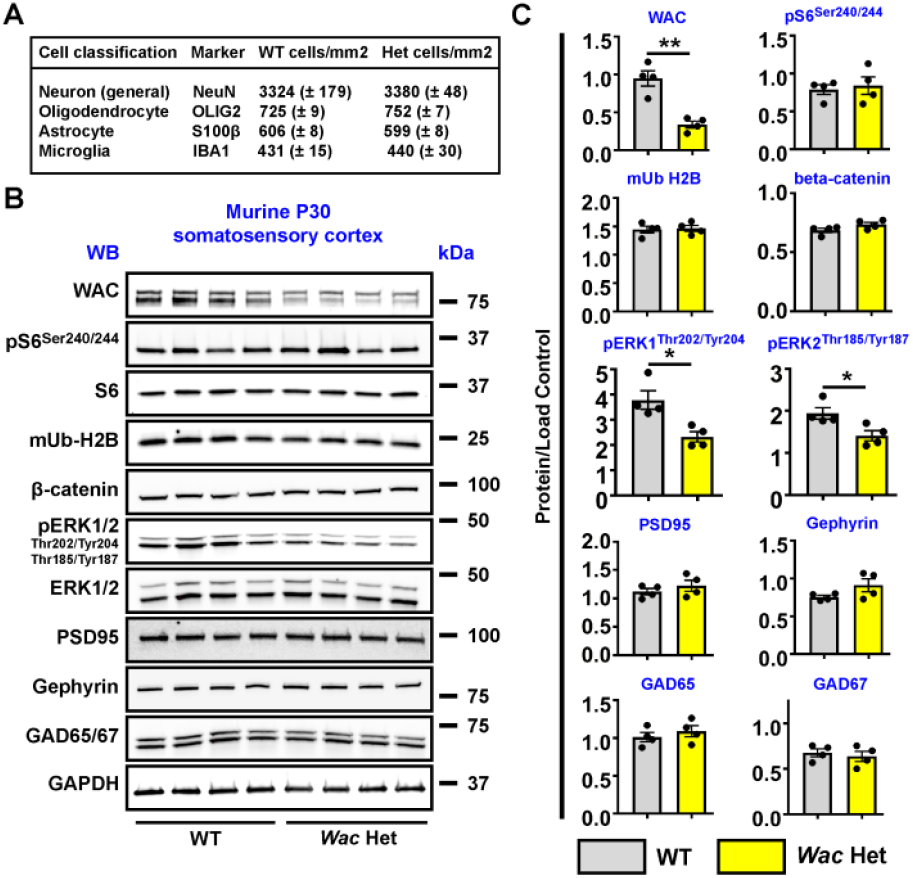
Cell counts and biochemical assessment of key markers in P30 *Wac* WT and Het mice. (A) WT and *Wac* Het somatosensory cortices were counted for major cell classes, including markers of neurons and glia, n = 3 each group, parentheses indicate Standard error of the mean. (B) P30 somatosensory cortex tissue was probed via western blotting for WAC and proteins known to be altered in other models or of brain markers. (C) Quantification of protein bands were normalized to GAPDH or the total protein if a phosphorylated protein. Data are expressed as the mean ± SEM, n = 4 all groups. * p < 0.05 and ** p < 0.01. Abbreviations: (WB) western blot and (kDa) kiloDaltons.

**Figure 6:**
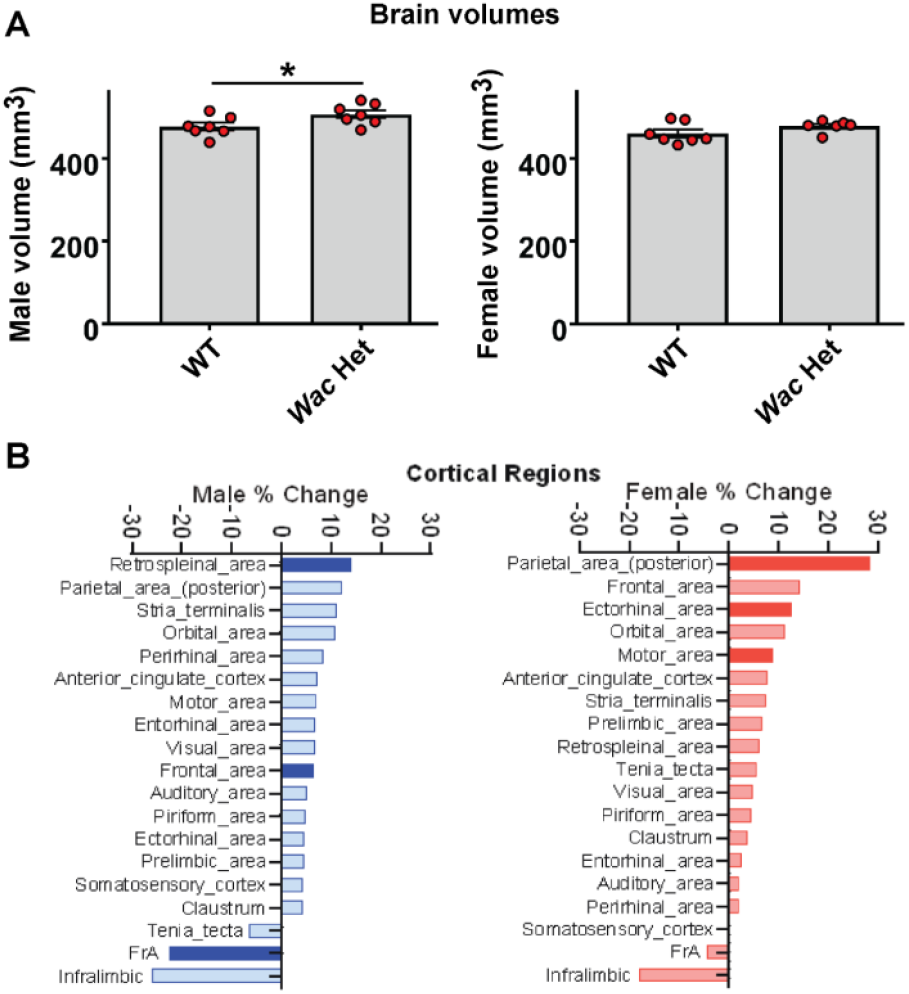
MRI reveals increased brain volume in mouse *Wac* Het males. (A) There was a significant increase in whole brain volumes in HET compared to WT male mice. Female HET mice did not exhibit significant changes in brain volume relative to WT. Data are expressed as the mean ± SEM; males and female WTs n = 7 and female Hets n = 6, *p < 0.05. (B) %change between cortical regions were larger in most HET mice areas, with distinct regions different between sexes. (Bolded bars indicate significant differences p < 0.05, t-test).

We also examined limbic regions and white matter track volumes in WT and *Wac* Het mice. As before, male mice had significant increases in both limbic and white matter volumes compared to females. Specifically, hippocampal ventral CA3 and dorsal CA1 regions were increased in males but not females (Figure S7, ventral CA3 male WT vs. male Het p = 0.0003 and dorsal CA1 male WT vs. male Het p = 0.04). Male specific increased volume was also observed in white matter dorsal fornix and corpus callosum tracts (Figure S8, dorsal fornix male WT vs. male Het p = 0.006 and corpus callosum male WT vs. male Het p = 0.01). As of this report, no DESSH syndrome individuals have been MRI assessed. However, these observations may be potential predictors of phenotypes that may arise as those with DESSH syndrome age into adulthood and whether there could be sex differences in humans.

### RNA sequencing of mouse forebrains reveals DESSH syndrome candidate transcripts

In a similar logic to our above tests, we chose to focus on the mouse model to assess potential RNA transcripts that are changed due to *Wac* loss of function since the mouse model may be more relevant to humans with DESSH. To discover potential molecular changes that could inform future mechanistic studies, we compared WT and *Wac* Het bulk RNA-seq at postnatal day 2 (P2), an age when mice are undergoing critical developmental milestones, including neuronal migration, cell death, axon and dendrite outgrowth as well as beginning synaptogenesis. We sequenced 14 WT and 10 *Wac* Het sex-balanced forebrain samples (Table S2), to a median sequencing depth of 50 million paired-end reads per sample (Figure S9; Table S2). As expected, we found significant correlations between sample sex and sequencing depth, and principal component analysis (PCA) dimensions; no outliers were detected in the PCA space (Figure S10; Table S2).

Differential expression (DE) analysis was performed using edgeR and included surrogate variable analysis (SVA) batch correction [31], which accounted for sex and batch biased transcriptomic variability. We found 7 and 16 genes were significantly upregulated and downregulated at FDR < 0.1, respectively; and 385 and 564 upregulated and downregulated genes were identified at a more inclusive P < 0.05 threshold (Figure 7A). DE was skewed towards downregulation, which is consistent with *Wac* function in promoting transcription [8]. While *Wac* gene expression was modestly upregulated in the mutants (log2FC = 0.09, P = 0.01, FDR = 0.5), coverage of floxed *Wac* exon 5 was reduced by 50% in HET samples (Figure S11), consistent with compensatory upregulation. As noted earlier, western blot found decreased WAC protein expression (Figure 5) and no evidence of mutant isoforms, suggesting that mutant *Wac* transcript is unlikely to be stably translated. Sex-stratified analysis identified 3 genes passing FDR < 0.1 (1 upregulated and 2 downregulated) in females, and 0 in males. At P < 0.05, the number of DE genes was higher in males, with 303 upregulated and 275 downregulated, in contrast to 193 upregulated and 149 downregulated genes in females (Figure 7A, Supplementary Tables 3-7); a volcano plot showing example up and down regulated genes is shown in Figure 7B.

**Figure 7:**
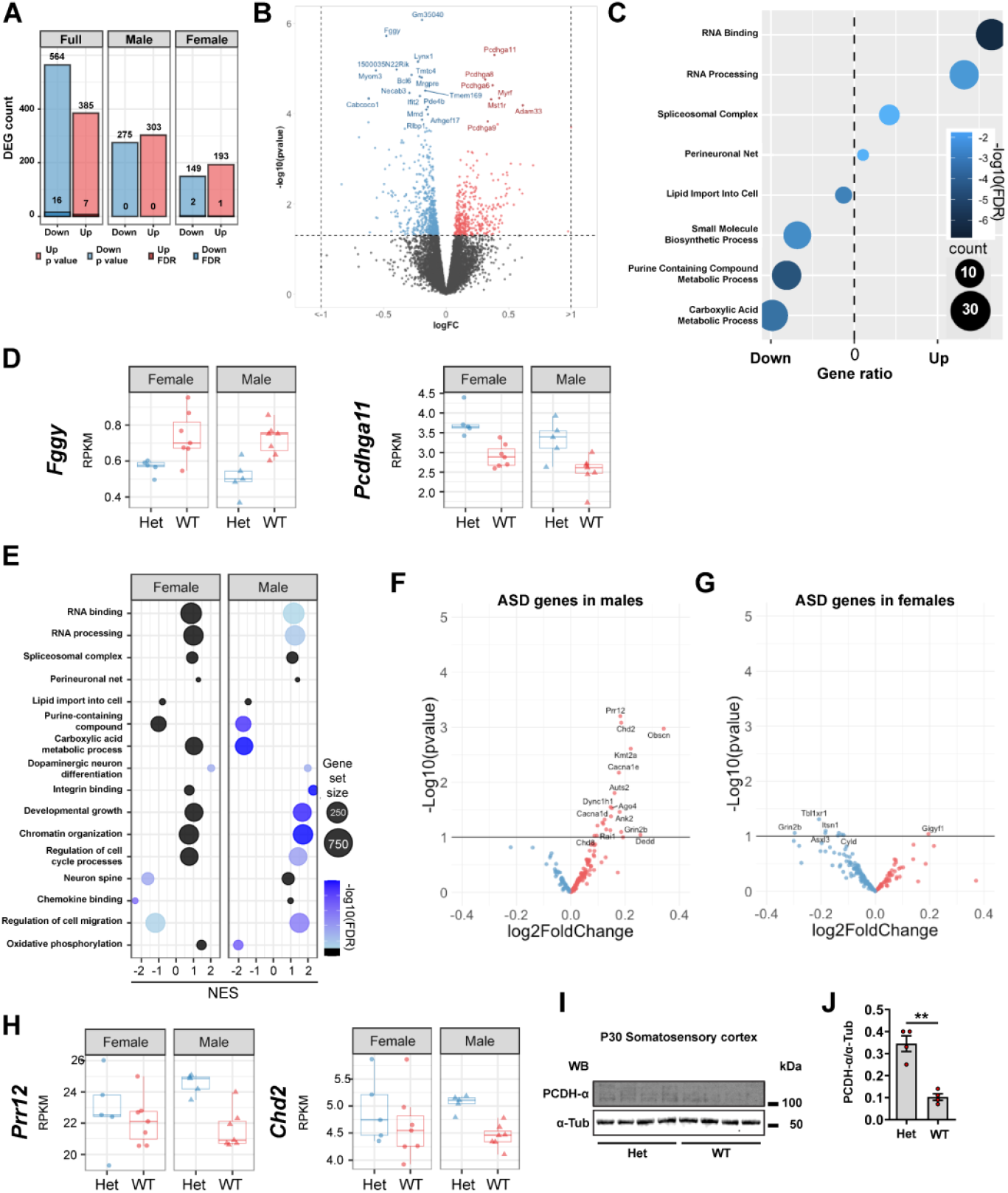
Transcriptomic characterization by bulk RNA-seq identified shared and divergent transcriptomic dysregulation in males and females. (A) Number of DE genes identified in the SVA-corrected DE analysis and sex-stratified DE analysis. Results are separated effect direction and significance (pvalue < 0.05 or FDR < 0.1). (B) Volcano plot representing DE of the SVA-corrected DE model in (A). (C) Selected GO analysis of significantly (P < 0.05) up and down-regulated genes in the SVA-corrected DE analysis. Results are colored by significance, the size indicated the number of DE genes belonging to that term and are ordered by the ratio of enrichment. (D) Examples of significant DE genes identified in the SVA-corrected model *Fggy* (FDR = 0.014) and *Pcdhga11* (FDR = 0.026), stratified by sex. (E) Curated GO terms from stratified GSEA analysis. Colors indicate significance, size the number of genes in the enriched term, and the enrichment score (NES). (F, G) Volcano plots showing the differential expression of ASD genes in males and females. (H) Examples of DE genes which show increased effect size and significance in males, *Prr12* (females, P = 0.99; males, P = 0.0006) and *Chd2* (females, P = 0.39; males, P = 0.0008). (I) Western blots showing PCDH-alpha abundance in WTs and Hets. (J) WB quantification comparing PCDH-alpha normalized to alpha-tubulin expression in P30 somatosensory cortices; n = 4 both genotypes and **p < 0.01. Data are expressed as the mean ± SEM.

We preformed gene ontology (GO) enrichment analysis of the DE genes passing a P < 0.05 threshold in the sex-combined model (Figure 7C). Upregulated DE genes were enriched for biological processes involved in RNA processing and splicing, whereas downregulated DEGs were enriched for metabolic processes clustered around purine and fatty acid metabolism. Gene set enrichment analysis (GSEA), which uses a rank-based approach to determine enriched pathways amongst up and down regulated genes, also identified RNA processing related terms to be upregulated and metabolic terms downregulated in *Wac* mutants (Figure S12A). *Fggy* is shown as an example of a metabolic gene with strong upregulation (Figure 7D). We also found a number of protocadherin genes (*Pcdhga6, Pcdhga8, Pcdhga9, and Pcdhga11*) were significantly upregulated (Figure 7B); *Pcdhga11* is shown which has consistent DE across sexes and samples (Figure 7D). We next tested for enrichment of disease associated genes using custom gene panels, finding no significant enrichment for high-confidence ASD genes based on rare mutations (https://doi.org/10.1038/s41588-022-01104-0) (n = 180) or the larger set of SFARI-associated ASD genes (https://doi.org/10.1186/2040-2392-4-36) (n = 1137), nor was there enrichment for genes associated with microcephaly (n = 1031), macrocephaly (n= 347), and epilepsy (n = 1318) (https://doi.org/10.1093/nar/gkz1021) (Figure S12B).

As DE analysis identified a potential sex bias with stronger effects in male *Wac* mutants (Figure 7A) and as the brain volume phenotype was larger in males (Figure 6A), we further investigated the sex-stratified DE results. Sex-stratified transcriptomic signatures were largely directionally correlated with each other, indicating most DEGs having a similar shift in RNA expression across sexes, with a large fraction of genes that had a stronger effect in males. (Figure S12C-E). We used GSEA to identify functional terms that were activated (upregulated) or suppressed (downregulated) in mutants (Figure 7E). Terms identified in the GO analysis of the full dataset showed concordant enrichment in both sexes (RNA binding, RNA processing, perineuronal net, lipid import into cells), but with higher significance in males. Notably, the metabolic related terms (purine-containing compound metabolic process, and carboxylic acid metabolic process) showed opposite directions with significant suppression in males. There was also concordant activation of dopaminergic neuron differentiation, integrin binding, developmental growth, chromatin organization, and regulation of cell cycle processes, with a stronger effect in males. Among the discordant signatures were suppression of neuron spine, chemokine binding and regulation of cell migration in females and suppression of oxidative phosphorylation in males. Using a permutation test to assess altered ASD genes, male DEGs (P < 0.05) showed a trend of enrichment for high-confidence ASD genes (P = 0.07) where females did not (P = 0.98) (data not shown). Examples of altered ASD transcripts are shown in Figures 7F, 7G, where males showed a unique alteration to several ASD genes compared to females. We highlight two ASD-associated genes with stronger effect in males versus females, *Prr12* (females, P = 0.99; males, P = 0.0006) and *Chd2* (females, P = 0.39; males, P = 0.0008) (Figure 7H). Finally, we wanted to validate if one of the more general changes between sexes in gene expression, that of the protocadherin alpha genes, was upregulated at the protein level. We thus probed P30 somatosensory cortices for pan protocadherin alpha via western blot (Figure 7I) as previously performed [32]; protein levels of protocadherin alpha were upregulated in the somatosensory cortices of *Wac* Hets (Figure 7J, p = 0.003).

RNA processing and splicing were identified as perturbed biological processes in the enrichment analysis and the RNF20/40/WAC complex has been reported to play a role in regulating chromatin accessibility and transcription through H2B ubiquitination [8], as well as mediating transcription elongation, and indirectly, splicing [8, 33]. We performed event-based differential splicing (DS) analysis using MAJIQ2.5 pipeline [34]. We found 13 DS junctions passing thresholds of 10% delta percent spliced in (dPSI) between WT and *Wac* Het, at 80% confidence level. As expected, we found two significant events representing floxed *Wac* exon skipping, and one intron retention in the Cre transgene locus Gt(ROSA)26Sor. We also detected skipped exons in *Snhg14*, alternative splice donors in *Dnmbp* and *Trmt61b*, alternative splicing acceptors in *Ssrp1* and *Trmt13*, retained introns in *Gm21992* and *Pigf*, and reduced intron retention in *Zc3h7a* (Figure S13, Supplementary Table 8). None of these genes were identified as significantly differentially expressed. Overall, results of DE analysis capture modest transcriptional differences with evidence for stronger DE effects in males, as well as some evidence for differences in splicing events in the *Wac* Het mutants. Overall, our data shed new light on the transcriptomic mechanisms and phenotypes of *Wac* haploinsufficiency and provide further context for the existing studies of DESSH syndrome.

## Discussion

DESSH syndrome is a monogenic disorder that results in variable degrees of developmental delay/intellectual disability, craniofacial dysmorphic features, ophthalmological and gastrointestinal problems, hypotonia and other neurological changes that may underlie the elevated co-diagnoses of ASD, ADHD and seizures. Previous work identified distinct molecular and cellular changes, including alteration to the mTOR signaling pathway and nuclear regulation of transcription that may also overlap with our studies. A major limitation is that these previous studies were restricted to invertebrates and cell lines but not vertebrates. Thus, we sought to test if genetic depletion models of the *Wac* gene in vertebrates could recapitulate some of the reported symptoms in humans and further use our mouse model to probe novel anatomical and molecular changes for future studies. While the mouse and zebrafish models mostly recapitulated most core DESSH symptoms tested herein they were not completely aligned. Importantly, mouse Hets recapitulated many DESSH symptoms while zebrafish *waca* KOs were needed to realize similar phenotypes. Since zebrafish have both *waca* and *wacb* genes, it is possible that some compensation occurs and only a complete deletion of one of the genes, here *waca*, could reveal symptoms that reflect DESSH patients.

While this study describes two novel vertebrate models for DESSH syndrome the mechanisms underlying our findings still need to be elucidated. This will be done in future studies using conditional *Wac* deletion models that are predicted to not result in lethality due to limited deletion of *Wac* in distinct brain or other tissues. It is possible that some mechanisms might be conserved from previous studies examining *Wac* in cell lines and drosophila [8, 10]. We previously found that the WAC protein still localizes to the nucleus and has a speckled pattern in mammalian neurons [7], similar to the first study examining *Wac* in a cell line [33]. Thus, a nuclear function for *Wac* is still highly likely and needs to be further tested. In addition, the WAC protein’s WW, coiled-coil and disorganized conserved domains are likely to mediate distinct adaptor complex properties and potential other regulatory functions that have yet to be uncovered.

Craniofacial dysmorphism and social behavior deficits are highly prevalent in DESSH, with craniofacial dysmorphism existing in almost 100% of those diagnosed with DESSH while autism behaviors are seen in roughly 30% of patients (Marwan Shinawi, personal communication), which were trending in mice and profound in zebrafish. We also found a robust sensitivity to a seizure inducing drug in mice but not zebrafish. Together, each model revealed unique strengths and better connections to core DESSH symptoms. Despite these differences, each model showed impacts upon GABAergic neurons. These changes in GABAergic neuron markers may or may not underlie the common behavioral social changes in each model while the seizures may have a different underlying mechanism. Regardless, these data present two novel vertebrate DESSH models that could provide new insights into *Wac* loss of function and the impacts that this gene regulates upon brain function over an evolutionary timeline.

We and others have observed a decrease of PV expression but no loss of MGE-lineage interneurons in other syndromic mouse models like *Cntnap2* and other models where just PV protein levels alone were assessed [25, 35, 36], suggesting this may be a common feature in some monogenic syndromes with overlapping phenotypic features. Individuals with DESSH syndrome can have seizures [1, 3, 4, 15] and we found that subthreshold GABAA receptor inhibition elicited seizures in *Wac* Het mice. The reaction to PTZ suggests that their brains are shifted towards increased excitability. It is also possible that this more excitable brain may influence the expression of PV, as we and others have found that brain activity can alter PV expression [27, 37]. Thus, it is still unclear whether the loss of PV leads to seizures in our model or if the elevated brain excitability could lead to changes in PV expression. No matter the cause, others have noted the importance of PV expression on behaviors relevant to ASD [38], suggesting that approaches to restore PV expression may be a future therapeutic to alleviate some symptoms of DESSH syndrome. Moreover, syndromes like DESSH may additionally benefit in the future from stem cell based therapies that supplement brains with inhibitory neurons, which is currently being tested to treat epilepsy and other conditions with an imbalance of inhibition [39, 40].

Some novel findings beyond phenotyping of our vertebrate models manifested that may be relevant for human symptoms. First, mutant male mice exhibit volumetric increases in several brain regions as adults, suggesting sex differences may arise in DESSH patients as they age and specific brain regions should be further explored. Zebrafish also have craniofacial extensions in the forebrain region. In addition, while only examined in mice thus far, transcripts changed in male Het mice encompassed more ASD risk genes, which may have future implications for understanding *Wac* as a regulator of other ASD risk genes in a sex-dependent manner.

Finally, our transcriptomic characterization of the *Wac* mouse model at P2 identified DE genes enriched in RNA processing and metabolism that may be future targets to understand underlying mechanisms. The magnitude of DE was stronger in males and although the signature was largely concordant in effect direction between the sexes, this could be indicative of a sex-based transcriptional response to reduced *Wac* dosage. Differential splicing signatures were comparatively more subtle, with 13 altered splicing events in 9 genes, including *snhg14*, an antisense transcript in the *Ube3a* locus, associated with Angelman and Prader-Willi syndrome. Of interest considering various sex-specific phenotypes associated with *Wac* haploinsufficiency and DESSH, male *Wac* mutants had stronger transcriptional phenotypes and a significant enrichment for ASD-linked genes among DEGs, suggesting possible convergence of neurodevelopmental pathways. Our forebrain analyses raise questions regarding the cell-type specific and developmental aspects of transcriptomic phenotypes, which could be addressed by single cell methods and longitudinal studies in the future. Overall, our transcriptomic results support roles for *Wac* during neurodevelopment and gene expression. We generated new vertebrate models to assess DESSH syndrome and their role during development in divergent vertebrate species. These phenotypes are a first attempt at understanding this complex ultrarare disorder and offer initial tools to probe further biological mechanisms and therapeutics.

## Materials and methods

### Animals

#### Mouse model

We thank the Wellcome Trust Sanger Institute Mouse Genetics Project (Sanger MGP) and its funders for providing the conditional mutant mouse gamete (C57BL/6N-Wac<tm2c(EUCOMM)Wtsi>/Wtsi). Funding information may be found at www.sanger.ac.uk/mouseportal, associated primary phenotypic information at www.mousephenotype.org and data about the development of these technologies [41–45]. Sperm harboring the *Wac* conditional allele was used to fertilize C57BL6/N donor eggs; progeny were then genotyped via polymerase chain reaction (PCR). We next bred *Beta actin-Cre* mice [12] with *Wac*^*Flox*^ mice to generate wild type (WT) and constitutive Het mice. After germline recombination, *Wac* Het mice were backcrossed and bred with CD-1 mice for at least three generations before being used for experiments. CD-1 outbred mice were used due to *Wac*’s importance for survival as these mice are more robust and can generate and care for more progeny; we attempted to collect embryonic constitutive KOs as early as E12.5 but never collected a KO, suggesting early embryonic lethality. All murine experiments were performed under the approval of Michigan State University’s Campus Animal Resources. ***Zebrafish model***. Zebrafish (*Danio rerio*) were maintained at 28.5°C with a 14 hours light/10 hours dark condition and fed three times a day. Zebrafish embryos were cultured in egg water (pH 7.4) and embryonic stages were determined by standard method [46]. All zebrafish experiments were approved by the Institutional Animal Care and Use Committees (IACUC) of Chungnam National University (CNU-00866) and zebrafish reared by standard lab protocols. For all assessments, males and females were tested. If no differences were noted, the data were combined for analyses. If differences were noted than the data were separated and presented independently. For all experiments both male and female animals were assessed but only reported as separate sexes if it became apparent that there were differences for specific phenotypes. Otherwise, all data were combined.

### Generation of *waca* KO zebrafish

Zebrafish *waca* gene sequence was acquired from the NCBI database (*waca*: NM_199660.1). The primer for in vitro transcription of sgRNA was designed by 5 ′ - TAATACGACTCACTATAGCTACTACAACTGCAGGACAGGTTTTAGAGCTAGAA - 3′ for the *waca* target site. Primer contained 5’ T7 promoter and 15 nucleotides at 3’ complement to the universal primer. Template for *in vitro* transcription of the sgRNA to knock-out *waca* was produced by PCR with primer 5’ – TAATACGACTCACTATAGGGAGGAAGGACTGGCACACGTTTTAGAGCTAGAA – 3’ [34]. *In vitro* transcription was carried out using PCR products and MaxiScript T7 Kit (Ambion). Cas9 expression vector pT3TS-nCas9n (Addgene), linearized with XbaI (NEB) and purified via DNA extraction kit (ELPIS). Cas9 mRNA was synthesized using mMESSAGE mMACHINE T3 Kit (Ambion) and tailed poly (A) with E. coli Poly (A) Polymerase (NEB). sgRNA of *waca* was injected with Cas9 mRNAs into 1-cell stage embryos.

### Behavior (mouse)

6-8 week aged mice of both sexes were tested; no differences were observed in any assay between sexes and data represent both groups. The persons performing the behaviors and analyses were blinded to the genotypes.

#### 3-chamber social interaction

During habituation, test mice had 10 minutes in the center of a rectangular chamber divided into 3-chambers by plexiglass containing small holes to navigate. Side chambers contained empty holders, with small holes allowing snout contact. Following initial habituation, an unfamiliar sex- and age-matched mouse (habituated to round holders/cages previously) was put in one of the chambers randomly with doors of the lateral chambers closed and a novel object in the other. The test subject explored the apparatus for 10 minutes. Sniffing time was recorded using an overhead video camera and analyzed using ANY-Maze software. Time spent in each chamber and interaction with mouse/object was recorded.

#### Y-maze

Individual mice were placed at the end of one arm of a Y-maze and allowed to explore for five minutes while being filmed by an overhead camera. Entries into all arms were noted (four paws need to be inside the arm for a valid entry) and a spontaneous alternation is counted if an animal enters three different arms consecutively. % of spontaneous alternation were calculated according to following formula: [(number of alternations) / (total number of arm entries − 2)] × 100.

### Behavior (zebrafish)

#### Novel tank assay

Novel tank assay was performed as previously described [47–49]. 3 month/older WT and *waca* KO sibling zebrafish were tested in the behavior tank (24 x 15 x 15 cm). The back and sides were covered with non-transparent white paper. Behavior tests were recorded for 20 minutes using a video camera (Sony, HDRCX190). All fish were returned to facility system tanks after completion of the behavioral tests. Recorded video files were analyzed using EthoVision XT7 software (Noldus).

#### Social cohesion test

Zebrafish group social cohesion was measured [47, 48]. During one session, 5 WT sibling or *waca* KO sibling zebrafish were placed in the behavior tank. Fish groups were recorded for 20 minutes using a video camera (Sony, HDRCX190). Videos were analyzed using 12 individual screenshots taken every 10 seconds for 3-5 minutes as early phase and 17-19 minutes as late phase. Distances between individual zebrafish in the group were measured using ImageJ for each screenshot and compiled.

### Cranio-facial analyses

#### Mouse model

The skulls were stained with Alizarin red for bone as previously published [50]. Quantification of the suture and fontanel areas and the skull width was performed from photographs using ImageJ. ***Zebrafish model***. Zebrafish cartilage was stained by Alcian Blue (Sigma) as previously described [51, 52]. Zebrafish larvae at 13dpf were fixed with 4% paraformaldehyde (PFA) overnight at 4ºC and treated with bleaching solution (3% H_2_O_2_/0.5% KOH in distilled water) for 30 minutes, rinsed out with Phosphate buffered saline with 0.1% Tween 20 (PBST). Specimens were transferred to methanol to store at −20º. Zebrafish cartilage was stained by Alcian blue solution (0.1% Alcian blue with 70% EtOH) for 1 hour and then dehydrated in 70% EtOH. Specimens were rinsed 100% PBST and then transferred to 90% glycerol with 1% KOH to analyze on a dissecting microscope.

### Genotyping

#### Mouse model

Primers to detect the recombined allele of the *Wac* genetic locus: Forward 5’-AGCTATGCGTGCTGTTGGG-3’ and Reverse 5’-CAAATCCCACAGTCCAATGC-3’. Thermocycling conditions were: 95ºC 3 minutes, (95ºC 30 seconds, 58ºC 30 seconds, and 72ºC 45 seconds for 35 cycles), 72ºC 3 minutes. Sanger sequencing to validate the recombined locus was performed by GeneWiz using the same primers on gel-purified PCR products. ***Zebrafish model***. Primers to detect the alleles of *waca*: Forward 5’-ATTTGAACCGGCAGATGATT-3’ and Reverse 5’-TCAGTGGGAGAAACCCAAAG-3’. Thermocycling conditions were: 95ºC 5 minutes, (95ºC 30 seconds, 60ºC 40 seconds, and 72ºC 20 seconds for 40 cycles), 72ºC 7 minutes and 15ºC 10 minutes. Primers to detect the genotypic alleles of *wacb*: Forward 5’-ATACCAGTCAAAGAGTCACTCAGCGAATGA-3’ and Reverse 5’-CACGTTCACCGGCCCCGTGAGAGAGACCAC-3’.

### Immuno-fluorescent staining

At P30, mice were transcardially perfused with phosphate-buffered saline (PBS), followed by 4% paraformaldehyde. The brains were then removed and postfixed in PFA for 30 minutes. Brains were transferred to 30% sucrose for cryoprotection overnight after fixation and then embedded in optimal cutting temperature compound before coronally sectioned at 25µm via cryostat. Sections were permeabilized in a wash of PBS with 0.3% Triton-X100, then blocked with the same solution containing 5% bovine serum albumin. Primary antibodies (details can be found in Supplementary Table 9) were applied overnight at 4ºC, followed by 3 washes. Secondary antibodies (Alexa-conjugated 488 or 594 anti-rabbit or mouse, 1:300, Thermo-Fisher) were applied for 1-2 hours at room temperature, followed by 3 washes and cover slipped with Vectashield. Two-three images from the somatosensory cortex were acquired that contained all cortical layers. Cell counts were performed on these images using the cell counter function in Image-J software and images from the same animal were averaged to report a biological “n”. Several hundred cells were counted per brain and 3-4 biological replicates performed for each marker.

### MRI acquisition and analysis

High-resolution ex vivo MRI was conducted on brains that were transcardially perfused with 4% paraformaldehyde, using a 9.4T Bruker Advance (Bruker Biospin, Billerica, MA, USA) with the following parameters: TR/TE 3397/10.6, 10 evenly spaced echoes 10ms apart, matrix 1.75 X 1.25 (91um isotropic), and 30 slices 0.5mm thick. T2-weighted imaging (T2WI) and T2 relaxation maps were computed as we previously described [53]. T2WI scans were skull stripped (masking) using the segmentation tool from the ITK-SNAP software (version 3.8.0, RRID:SCR_002010) [54]. The T2WI mask was then used to generate a 3D reconstruction of the brain. T2WI parametric maps were generated and corrected for bias field inhomogeneities prior to registration. From T2WI data, brain and regional volumes and T2 relaxation times were obtained. The Australian Mouse Brain Mapping Consortium (AMBMC) atlas [55, 56] was utilized to extract 40 bilateral regions using non-linear registration to each subject’s T2 images. Regional labels were applied using Advanced Normalization Tools (ANTs) and manually inspected for registration quality. Outliers were identified by calculating the interquartile range (IQR) for each region where 1.5IQR above the first quartile or below the third quartile were identified and excluded using Microsoft Excel.

### Microscopy

Mouse fluorescent images were acquired using a Leica DM2000 microscope with a mounted monochrome DFC3000G camera. Fluorescent images were adjusted for brightness/contrast and merged using Image-J software. The skulls were photographed with Nikon SMZ1500 stereomicroscope and Nikon DSRi1 camera. Zebrafish images were acquired using Leica TL 5000 microscope with DFC 7000T camera.

### PTZ induced seizure assessments

1 gram of Pentylenetetrazole (PTZ) powder was dissolved into 250 ml of PBS and injected intraperitoneally at 50 mg of PTZ per gram of mouse body weight. After injection, mice explored a clean cage for 20 minutes. Seizure severity was scored based on a modified Racine scale [57]: 1) Freezing or idle behavior; 2) Muscle twitches and/or rhythmic head nodding; 3) Tail arching over the mouse’s backside accompanied by the mouse assuming a hunched posture; 4) Forelimb clonus; 5) Tonic-clonic seizures with a recovery to normal behavior; 6) Uncontrolled running/jumping/hyperactivity; 7) Full body extension of limbs; 8) Death.

### RNA-Sequencing and bioinformatics analysis

Twenty-four mouse brains (14 WT and 10 *Wac* HETs; 7, 7 and 5, 5, males, females, respectively) were collected at P2 following instant decapitation. Samples were snap frozen on dry ice and stored at −80C until RNA preparation. On the day of RNA isolation, left and right forebrain hemispheres were separated from the hindbrain and the total RNA was consistently isolated from the right forebrain hemisphere. Total RNA was obtained using Ambion RNAqueous Total RNA Isolation Kit (cat# AM1912) and assayed via Agilent RNA 6000 Nano Bioanalyzer kit/instrument. Sample RIN scores ranged from 6.9 to 9.8, with a mean RIN score of 9.4.

Poly-A-enriched mRNA libraries were prepared at Novogene using Illumina reagents. Libraries were sequenced using Illumina NovaSeq 6000 S4 system, paired-end 150 (PE150) method. Reads were aligned to mouse genome (GRCm38/mm10) using STAR (version 2.5.4b) [58], and gene counts were produced using featureCounts [59]. On average, 50 million paired-end reads (PE) aligned per sample, with a range of 32 to 84 million reads.

Data quality was assessed using FastQC. FastQC: A Quality Control Tool for High Throughput Sequence Data. Available online at: http://www.bioinformatics.babraham.ac.uk/projects/fastqc/ and principal component analysis (PCA) was used to determine presence of sample outliers. All 24 samples were qualified for the analysis. Raw RNA-seq fastq files and a gene count matrix is available on GEO (GSE264597).

Bioinformatic analysis was performed using R programming language version 4.2.1. R: A language and environment for statistical computing. R Foundation for Statistical Computing, Vienna, Austria. URL https://www.R-project.org/ and RStudio integrated development environment version 2023.06.0. RStudio: Integrated Development Environment for R. Posit Software, PBC, Boston, MA. URL http://www.posit.co/. Plots were generated using ggplot2 R package version 3.4.0, URL: https://link.springer.com/book/10.1007/978-3-319-24277-4. Heatmaps were generated using pheatmap R package 1.0.12.

### Differential expression (DE) analysis

For DE analysis we used edgeR R package [60]. Genes with a minimum of 1 counts per million (CPM) in at least six samples were included in the analysis. The first surrogate variable from the SVA-batch-correction method was used as a covariate in the edgeR GLM model [61]. For sex-stratified DE, we used a threshold of CPM > 1 in at least 3 samples, and no batch correction. Reads Per Kilobase per Million mapped reads (RPKM) were used for plotting summary heatmaps and expression data of individual genes.

### Gene ontology enrichment analysis

To test for enrichment of GO terms we used the clusterProfiler R package version 4.14.6 (Bioconductor). Mouse Gene Ontology (GO) data for biological processes (BP), molecular function (MF) and cellular component (CC) were downloaded from Bioconductor (org.Mm.eg.db). For the analysis presented here, DE genes with a P value < 0.05 were used as input, and we required a minimum set of 5 and maximum of 1,000 as well as at least 2 significant DE genes in a GO term. We a Benjamini-Hochberg (BH) correction on significance, a strategy recommended for gene set analysis that accounts for multiple testing comparisons. We reported terms with p-value<0.05. The set of DE genes was compared against the background set of genes expressed in our study based on minimum read-count cutoffs described above. To test for enrichment of GO terms we used the fgsea R package version 1.32.4, via clusterProfiler. To control for transcriptional noise, we only used protein coding transcripts with a logCPM > 1. We tested against terms for BP, MF, and CC using a minimal gene set size of 5 and maximum of 1,000. We again used a Benjamini-Hochberg (BH) correction on significance and reported terms with P < 0.05. For the custom GSEA we pulled genes associated with epilepsy (C0016399), microcephaly (C1837501), and macrocephaly (C1836599) from DisGeNET (https://doi.org/10.1093/nar/gkz1021), SFARI genes from (https://doi.org/10.1186/2040-2392-4-36), and high-confidence ASD genes from (https://doi.org/10.1038/s41588-022-01104-0) with a FDR < 0.05.

### Permutation test

To determine significance of the overlap between DE genes identified in RNA-seq and disease gene sets, we used a permutation test. We used the DE genes with P < 0.05 to compare against the ASD gene set from Fu et al (https://doi.org/10.1038/s41588-022-01104-0) FDR < 0.05 and the epilepsy gene set from DisGeNET (https://doi.org/10.1093/nar/gkz1021). To generate a null distribution, we performed 10,000 permutations. In each iteration, a random set of genes equal in size to the DE gene list was sampled without replacement from the background gene set of those tested in DE analysis. The overlap between the disease gene set and each permuted gene set was recorded. And a p-value was calculated as the proportion of permutations in which the overlap was greater than or equal to the observed overlap.

### Analysis code is available at

https://urldefense.com/v3/_ https:/github.com/NordNeurogenomicsLab/Publications/tree/master/Lee_et.al._2024*2C_Complimentary_vertebrate_Wac_models_exhibit_phenotypes_relevant_to_DeSanto-Shinawi_Syndrome__;JQ!!HXCxUKc!0tNGmjep5OYi3Z-2ICpXdVi5iivlrNqffntUIxCwJDU1JgPyTtFTU7qg0XsXJuzFU4f_7M5FVXwiPayDnAJaasqO$

### Western blots

P30 somatosensory cortices were dissected and frozen on dry ice. Next, they were lysed in RIPA buffer containing protease and phosphatase inhibitors and combined with Laemmli buffer containing 2-Mercaptoethanol and incubated at 95ºC for 5 minutes to denature the proteins. Equal amounts of protein lysates were separated on 10% SDS-PAGE gels and then transferred to nitrocellulose membranes. The membranes were washed in Tris-buffered saline with Tween-20 (TBST) and then blocked for 1 hour in TBST containing 5% non-fat dry milk (blotto, sc-2324 SantaCruz biotechnology). Membranes were then incubated with primary antibodies overnight at 4ºC, washed 3 times with TBST, incubated with secondary antibodies for 1 hour at room temperature and then washed 3 more times with TBST. Membranes were next incubated in ECL solution (BioRad Clarity substrate 1705061) for 5 minutes and chemiluminescent images obtained using a BioRad Chemidoc™ MP imaging system. Primary antibody details can be found in Supplementary Table 9, secondary HRP antibodies were used at 1:4000 (BioRad). Uncropped western blots are shown in Figure S14.

### Whole-mount *in situ* hybridization (WISH)

WISH was performed using DIG-labeled antisense RNA probes for *gad1b*. RNA probes were synthesized using DIG-RNA labeling kit (Roche) [62]. Zebrafish larvae were fixed in 4% PFA in PBS overnight at 4ºC then dehydrated with 100% Methanol. Larvae were rehydrated with 0.1% Diethyl pyrocarbonate with PBST (DEPC-PBST, Sigma Aldrich). Larvae were permeabilized with 10μg/ml proteinase K (Roche) according to developmental stage. Larvae were re-fixed for 15 minutes in 4% PFA, washed with DEPC-treated PBST, and pre-hybridized in HYB solution (50% formamide, 5X saline sodium citrate, 50μ g/ml heparin, 500 µg/ml torula RNA, 46mM citric acid pH 6.0, 0.1% Tween20) for 1 hour at 70ºC. Antisense DIG-labeled RNA probes were added in HYB solution and incubated for overnight at 70ºC. Larvae were washed in a preheated mixture of 50% saline sodium citrate containing 0.1% Tween-20 and 50% hybridization solution at 70ºC. Larvae were blocked and then incubated with 1/4000 concentration of anti-DIG Fab fragment conjugated with alkaline phosphatase (Roche) overnight at 4ºC. Larvae were washed in PBST then incubated in staining solution including NBT/BCIP as alkaline phosphatase substrate in the dark until sufficient staining appeared. Larvae were mounted in 90% glycerol in PBST and were visualized via dissecting microscope.

### Quantification and statistical analysis

Statistical analyses were performed using Prism versions 7 and 10, a p value of < 0.05 was considered significant. For parametric measures of two groups, a two-tailed T-test was performed; parametric measures of three or more groups utilized a ONE-Way ANOVA with Tukey post-test. For the male retropleinal cortex, male parietal cortex and female corpus callosum MRI data, the data were not distributed normally and we used Mann Whitney test. If other statistical analyses were used they are denoted in the results or figure legends.

## Acknowledgements

**AMS, DV, KU, DP-C, TEJ** and **XL** were funded by the Michigan State University/Corewell Health Corporation. **DP-C** was additionally supported by an Integrative Pharmacological Sciences Training Program (IPSTP) grant, 5T32GM142521. We would like to thank DubDub and Harrison Levon-James Allen for critical insights and providing resources during this study; **MP-V** and **JJ** were funded by NIDCR R01 DE026798; **KU, TEJ** and **XL** were also funded by the Cystic Fibrosis Foundation LI19XX0, the Cystic Fibrosis Research Inc., R01 HL153165-01A1. **ASN** was funded by NIMH R01 MH120513. **C-H K** was supported by grants from the National Research Foundation of Korea (RS-2024-00443043, RS-2024-00349650) and the Korea National Institute of Health (KNIH) research project (2026-ER0602-00, 2026-ER0807-00). RNA-seq data were funded by a Michigan State University Neuroscience Program grant. We would also like to thank Tom and Cathy Mall for their generous gift to support our research underlying autism genetics at Michigan State University.

## Conflict of interests

The authors report that they have no conflict of interests.

## Notes

### Competing Interest Statement

The authors have declared no competing interest.

### Summary of Updates

Final edits for an eLife revison: includes author addition and financial updates.

