## Supplemental main file for "Complementary vertebrate *Wac* models exhibit phenotypes relevant to DeSanto-Shinawi Syndrome" (from another version of this preprint: version 4, 2026-04-12)

**Supplemental material**

### Methods

Antibody labeling: Immuno-fluorescent staining and imaging of P30 mouse brain tissue was performed as described in the main text. Antibodies used: rat anti-CTIP2 (1:500, ab18465 Abcam), mouse anti-SATB2 (1:500, ab51502 Abcam) and rabbit anti-TBR1 (1:500, ab31940 Abcam).

Differential splicing (DS) analysis: Differential splicing (DS) analysis was performed using MAJIQ v2.5.1 (Vaquero-Garcia et al. 2023, PMID: 36869033). In short, the *majiq build* module was executed on WT (n = 14) and Wac Het (n=10) BAM files using GRCm38.95.gff3 genomic reference. DS quantification was performed using *majiq deltapsi* module, with two.majiq file groups delineating WT and Het samples. DS splicing was visualized using the *voila view* command with abs(E(dPSI)) threshold set to 0.1 and confidence threshold set to 0.8. Output files from the MAJIQ pipeline are deposited at Zenodo https://zenodo.org/records/10999584 with the accession number 10.5281/zenodo.10999584.

Electrophysiology: Coronal cortical slices (300 µm thick) were prepared (between postnatal ages 46 and 55 using methods previously described [1, 2]. Briefly, mice were deeply anesthetized with isoflurane before decapitation. Brains were then quickly removed and placed in a cold (~4°C) oxygenated slicing solution (95% O2, 5% CO2) containing (in mM): 3 KCl, 1.25 NaH2PO4, 10 MgSO4, 0.5 CaCl2, 26 NaHCO3, 10 glucose, and 234 sucrose. Slices were cut using a vibrating tissue microtome (Leica VT1200S) and then transferred into a chamber containing warm (32°C) oxygenated (95% O2, 5% CO2) artificial cerebrospinal fluid (ACSF) containing (in mM): 126 NaCl, 3 KCl, 1.25 NaH2PO4, 2 MgSO4, 2 CaCl2, 26 NaHCO3, and 10 glucose. Slices were kept at 32°C for 20 min followed by room temperature for an additional 40 min before recording.

For recordings, individual slices were transferred to a submersion recording chamber and continually perfused (~3 ml/min) with warm (32°C) oxygenated (95% O2, 5% CO2) ACSF containing (in mM): 126 NaCl, 3 KCl, 1.25 NaH2PO4, 2 MgSO4, 2 CaCl2, 26 NaHCO3, and 10 glucose. Neurons were visualized using infrared differential interference contrast (IR-DIC) and fluorescence imaging using a Zeiss Axio Examiner.A1 microscope mounted with a video camera (Olympus XM10-IR) and a 40x water-immersion objective. Whole-cell recordings were obtained from pyramidal neurons in layer 2/3 of the primary somatosensory cortex using borosilicate glass pipettes (4–6 MΩ tip resistance) containing a potassium-based internal solution (mM): 130 K-gluconate, 4 KCl, 2 NaCl, 10 HEPES, 0.2 EGTA, 4 ATP-Mg, 0.3 GTP-Tris and 14 phosphocreatine-K (pH 7.25, 290 mOsm). All whole-cell recordings were corrected for a 14-mV liquid junction potential.

Data were recorded and digitized at 20 or 50 kHz using Molecular Devices hardware and software (MultiClamp 700B amplifier, Digidata 1550B4, and pClamp11.1). Signals were low-pass filtered at 10 kHz before digitizing. During the recordings, the pipette capacitances were neutralized and series resistances (typically between 10 and 25 MΩ) were compensated online (100% for current-clamp, 70% for voltage-clamp). Series resistances were continually monitored throughout the recordings.

Analysis of electrophysiological data was performed in Molecular Devices and Microsoft Excel as described (Crandall et al., 2017). Resting membrane potentials (RMP, mV) were measured after break-in with no applied current. Input resistance (Rin, MΩ) was measured using Ohm's law by measuring the voltage response from rest to an injection of a small negative current (15-25 pA). Membrane time constants (τm, ms) were measured from the average response (see Rin above) by fitting a single exponential to the initial falling phase of the response (100–300 ms; omitting the 1st ms). Membrane capacitance (Cin, pF) was calculated by τm/Rin. Rheobase currents (pA) were defined as the minimum positive current (5 pA steps) to elicit an AP from a holding potential of −84 mV. AP properties were measured from the first spike evoked by the rheobase current. AP threshold (mV) was defined as the membrane potential at which its first derivative (dV/dt) exceeded 10 mV/ms. AP amplitudes (mV) were defined as the voltage difference between the threshold and the peak of the AP. AP half-widths (ms) were measured at the half-height between the threshold and the AP peak. The max rate of rise and decay (mV/ms) was defined as the maximal dV/dt during the rising phase and falling phase of the same AP, respectively. Frequency-intensity (F/I) relationships were obtained by holding the soma at −84 mV with intracellular current and injecting suprathreshold positive current (50 pA steps, 500 ms duration). F/I slopes

(Hz/pA) were determined using the initial frequency (reciprocal of the first interspike interval) over the entire F-I plot.

Spontaneous excitatory postsynaptic currents (sEPSCs) were recorded at -94 mV. Initially, individual events were manually selected and compiled to create an averaged sEPSC template that was then used to automatically detect similar events. Using this template, individual events were automatically selected by the software, but each event was also checked visually to confirm quality. 50 sEPCs were selected for each neuron for analysis of synaptic properties, including current amplitude and frequency. sEPSC amplitude and frequency were measured from events and automatically selected using our template. For each preselected event, the peak inward current amplitude was detected by the waveform template and was compared to a baseline immediately preceding each event.

<u>Mouse behavior</u>:

*Elevated plus maze*: Individual mice were placed in the center of a two open and two enclosed arm 50 cm height platform, facing an open arm [3]. Mice were allowed to explore for 5 minutes while being tracked and later analyzed by ANY-maze software. The % open-arm entries [(total number of entries/number of open-arm entries) × 100] and % time spent in the open arms were assessed.

*Open field*: Individual mice were placed in an open chamber with a center region designated by a 10 × 10 $cm^2$ area and the remaining area to the wall designated as periphery [4]. Mice roamed freely for ten minutes while being tracked by an overhead camera. The total distance travelled, and the mean speed, were analyzed using ANY-maze software as well as time spent in center versus periphery.

*Radial Arm Water Maze*: Individual mice were placed in a corner of a circular swimming pool containing 6 swim arms radiating from a central area [5]; water temperature of 26ºC. A hidden escape platform located at the end of one arm; mice were allowed up to 60 seconds/trial (15 trials total) on day 1 to find the platform in a room with 4 distinct spatial cues mounted onto each of the 4 walls of the room. Each trial was followed by a 15 second rest period on the platform and each trial will alternate having a visual cue on the platform and being hidden. If a mouse did not find the platform within 60 seconds, it was guided there and allowed the rest period. After learning trials, each mouse was dried with a small towel and allowed to rest for 30 minutes in a dry cage with a recirculating water blanket under half of the cage to provide warmth. On the second day, each mouse underwent 15 trials using only the spatial cues (hidden platform) and the number of errors (wrong arm entries) recorded by an overhead camera and reported.

*Whole-mount in situ hybridization (WISH)* procedures are described in the main methods.

**Supplementary Figure 1**

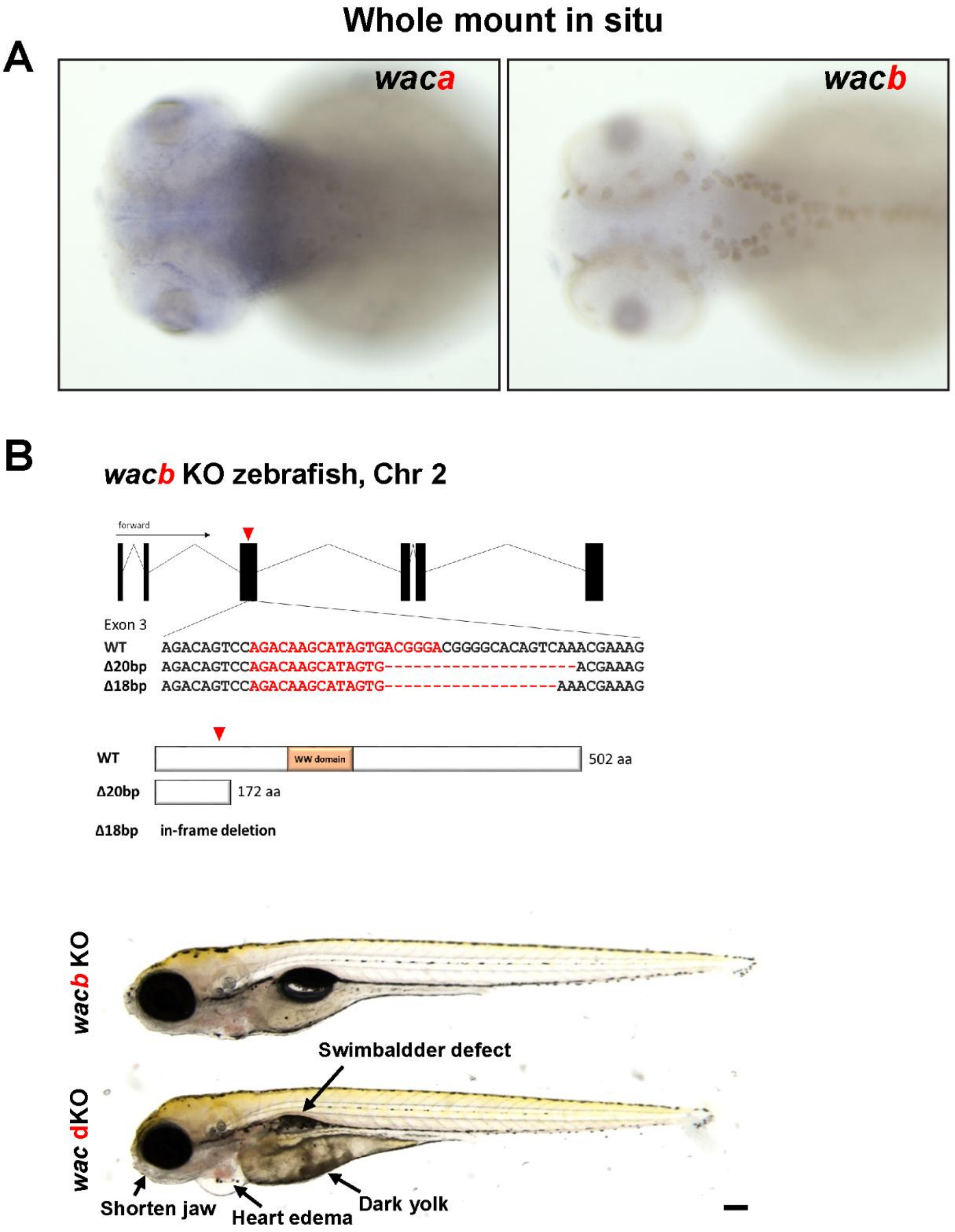


***waca* and *wacb* mRNA expression and generation of other zebrafish *wac* models**. (A) Whole-mount *in situ* hybridization analysis of *waca* and *wacb* mRNA in zebrafish at 72 hours post fertilization. *wacb* KO zebrafish were generated by CRISPR/Cas9; double KOs were generated by intercrossing *waca* and *wacb* zebrafish. KO target site for sgRNA is indicated on the gene structure (arrowhead). Predicted protein structures show premature truncation caused by frameshift deletion mutations (-20 bp for *wacb* KO). b) Morphological characterization of *wacb* KO and *waca,b* dKOs at 5 dpf. *wacb KO* zebrafish showed no obvious phenotypes at any stage analyzed, while *waca,wacb* dKO showed heart edema, dark yolk, and swimbladder defect, in addition to shortened jaw. *waca,wacb* dKO zebrafish did not survive over 7 dpf. Scale bar in (B) = 200 μm.

**Supplementary Figure 2**

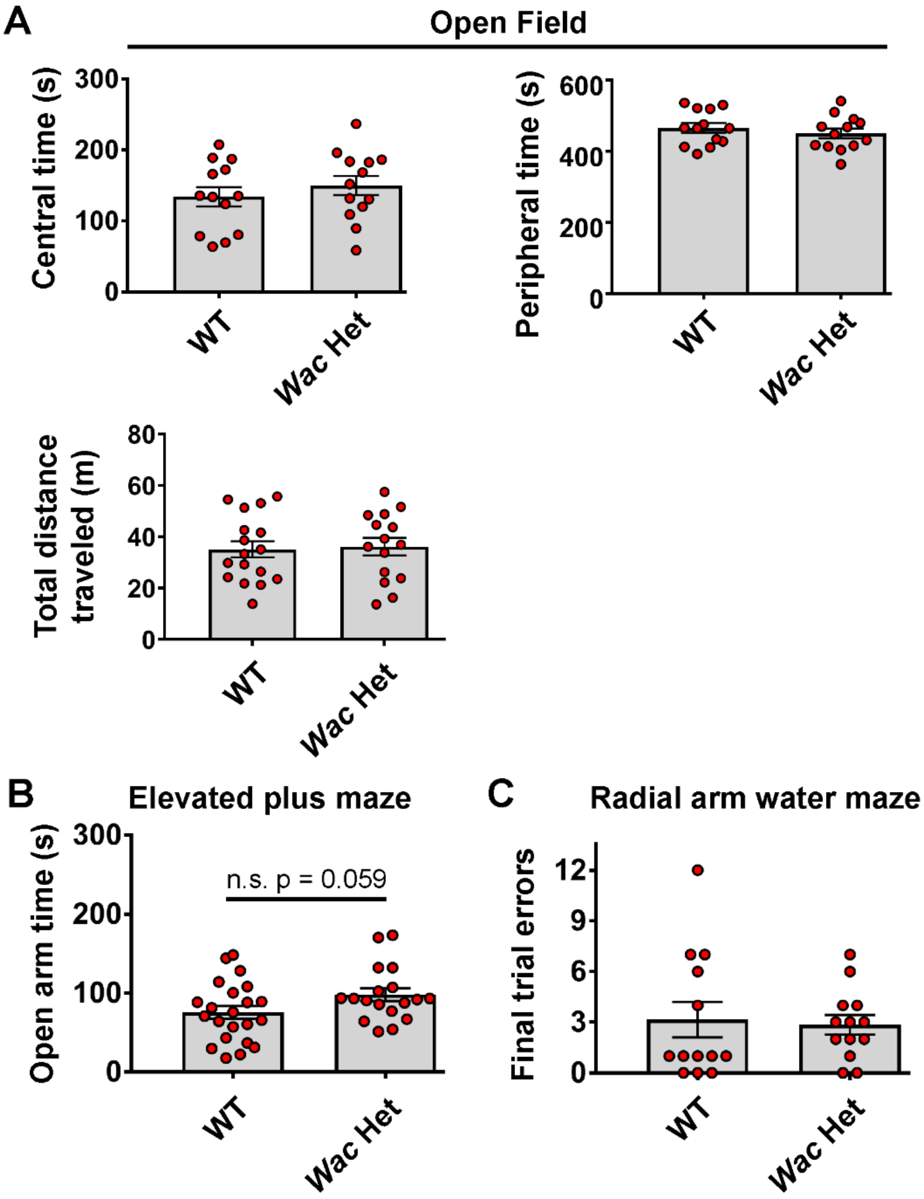


**Ambulatory, spatial memory and anxiety relevant tests in *Wac* Het mice**. Adult WT and *Wac* Het mice were assessed for ambulatory activity in the open field test (A); no changes were found in time spent in the center (top left graph) or periphery (top right graph), as well as total distance travelled (bottom left graph) between groups. (B) The elevated plus maze was used to determine whether mice preferred a safe enclosed arm or open arms. While not significant, *Wac* Het mice trended towards preference for the open arms. (C) A spatial memory test was used to test if *Wac* Hets had deficits in spatial memory but as a group no significant differences were observed. Data are expressed as the mean ± SEM, n = 13 both groups for open field periphery measurements, n = 17 (WT) and 15 (Het) for total distance. For radial arm water maze, n = 22 WTs and 18 Hets for Elevated plus maze.

### Supplementary Figure 3

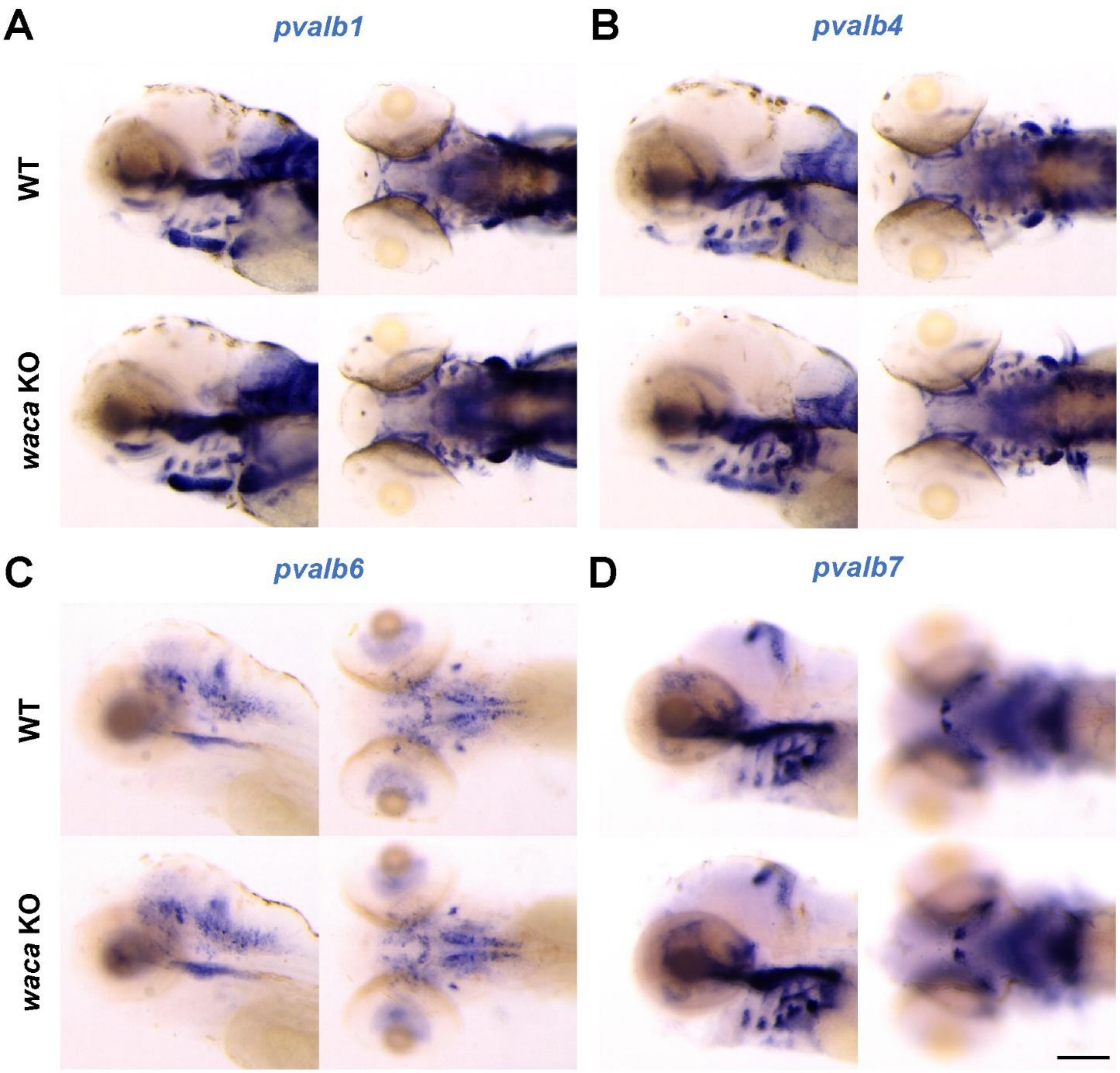


**Expression pattern of parvalbumin genes (*pvalb1*, *pvalb4*, *pvalb6*, and *pvalb7*) in *waca* KO zebrafish at 7 dpf**. (A-D) Lateral view in left panels; dorsal view in right panels; anterior is to the left. No significant change was detected between WT and *waca* KO. n = 7 for WT and n = 6 for *waca* KO for *pvalb1*. n = 5 for WT and n = 9 for *waca* KO for *pvalb4*. n = 4 for WT and n = 6 for *waca* KO for *pvalb6*. n = 7 for WT and n = 3 for *waca* KO for *pvalb7*. Scale bar in (D) = 200 μm.

**Supplementary Figure 4**

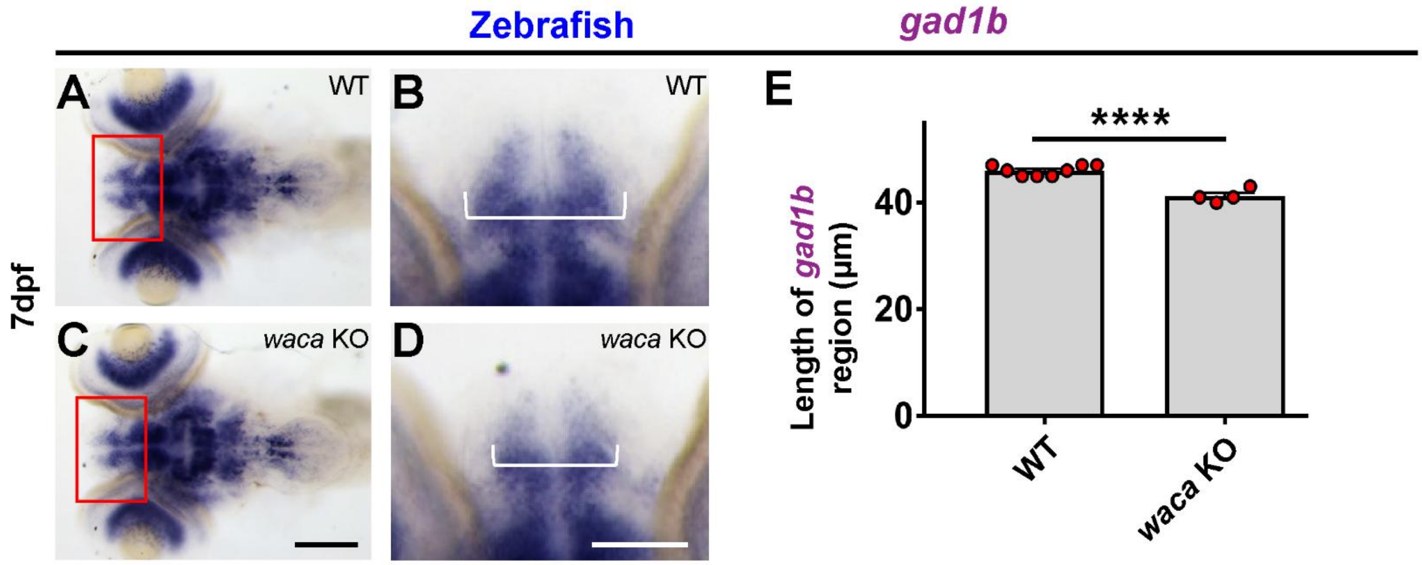


**Expression pattern of the *gad1b* gene in *waca* KO zebrafish at 7 dpf**. (A-D) Dorsal view of *gad1b* labeling, with anterior to the left (in left panels) and to the top (in right panels). Boxes highlight forebrain regions. Brackets demarcate area of *gad1b* expression. (E) Quantification of the *gad1b* area of expression between WT and *waca* KOs. Data are expressed as the mean ± SEM, n = 8 for WT and n = 4 for *waca* KO, ****p < 0.0001. Scale bars in (C) = 200 µm, (D) = 200 µm.

**Supplementary Figure 5**

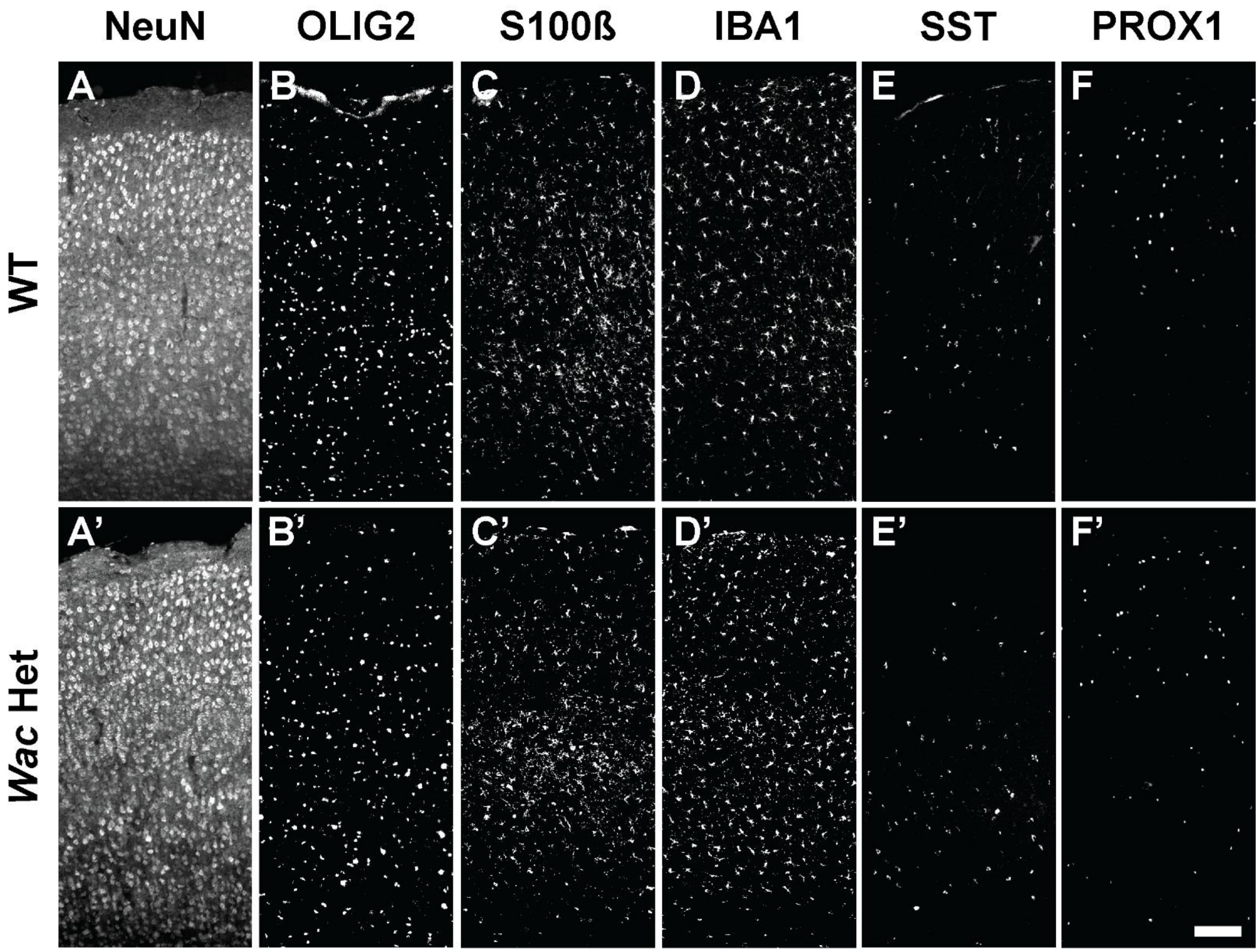


**Example images of cortical glial and neuron cell types in WT and *Wac* Het brains**. Scale bar in F’ = 100μm.

**Supplementary Figure 6**

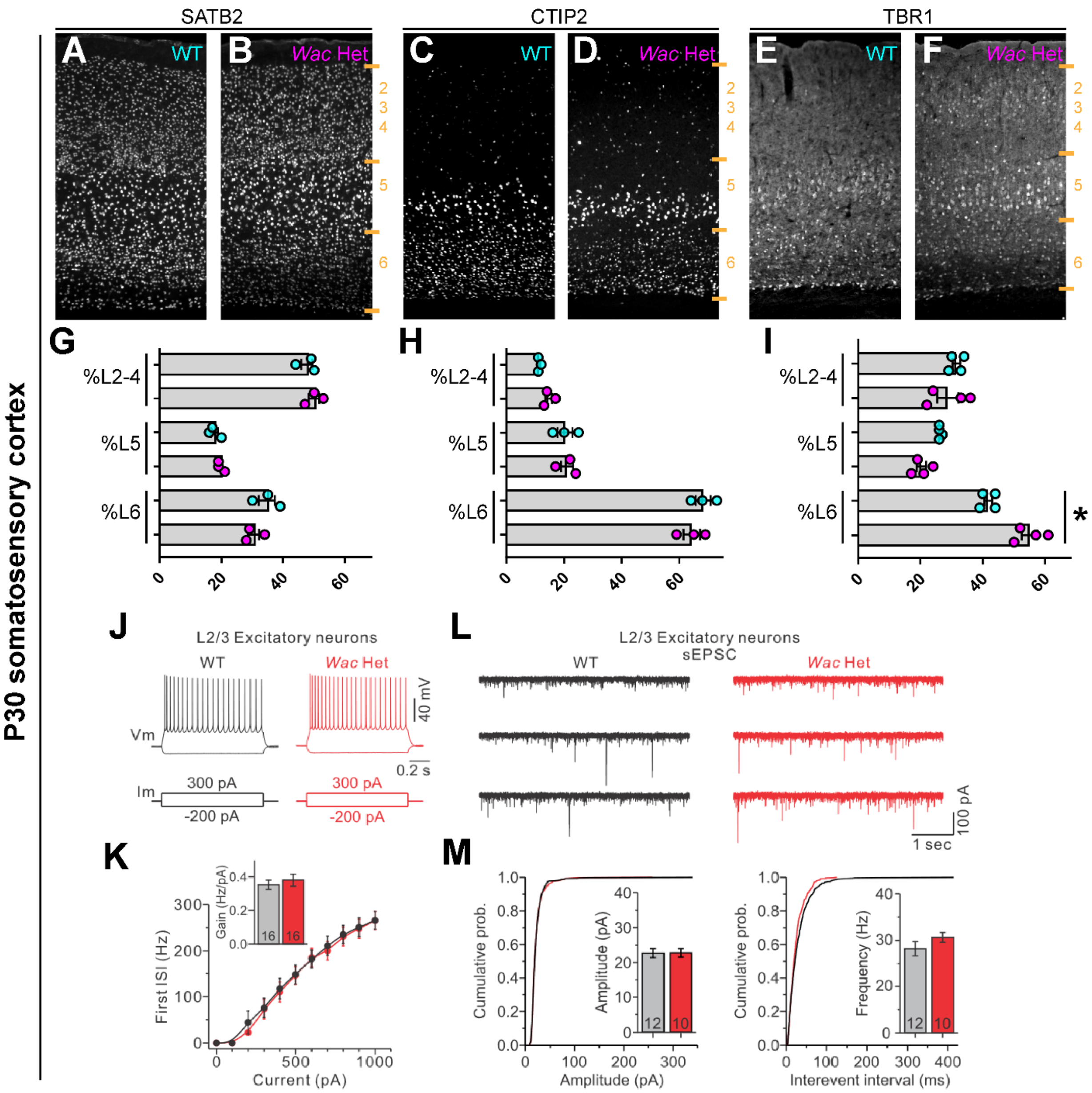


**Normal pyramidal neuron lamination, intrinsic membrane properties and spontaneous excitatory postsynaptic currents in the somatosensory cortex.** Representative images of pyramidal neuron laminar markers in somatosensory cortices of P30 WT and *Wac* Hets for SATB2 (A, B), CTIP2 (C, D) and TBR1 (E, F). Quantification of the % marker per layer (L) for SATB2 (G) n = 3, CTIP2 (H) n = 3 and TBR1 (I) n = 4. (J) Example voltage responses of wild-type (WT) and *Wac* Het L2/3 excitatory neurons to injected current. (K) Frequency-intensity (FI) plots showing the first interspike interval (ISI) frequency for WT and *Wac* Het L2/3 excitatory neurons following positive current steps (0-1000 pA; 100 pA steps; 1000 ms duration) (WT: 16 cells from 3 mice; *Wac* Het: 16 cells from 3 mice). Bar graphs show that the mean linear slopes (gain) of the FI curves were not significantly different (WT: 0.35 ± 0.03 Hz/pA; *Wac* Het: 0.38 ± 0.04 Hz/pA; p = 0.54, two-sample t-test). (L) Three representative traces of spontaneous excitatory postsynaptic currents (sEPSCs: 5 s per trace) recorded in voltage-clamp (-94 mV) from a WT and *Wac* Het L2/3 excitatory neuron. (M) Cumulative probability plots of sEPSC amplitudes (left) and interevent intervals (right) from WT and *Wac* Het L2/3 excitatory neurons (WT: 540 events, 12 cells from 3 mice; *Wac* Het: 450 events, 10 cells from 3 mice). Bar graphs show no significant differences in the mean sEPSC amplitude (left) or frequency when averaged from individual L2/3 excitatory neurons (WT amplitude: 22.7 ± 1.3 pA; *Wac* Het amplitude: 22.8 ± 1.2 pA, p = 0.94, two-sample t-test) (WT frequency: 36.4 ± 3.1 Hz; Wac Het: 41.3 ± 2.1 Hz, p = 0.23, two-sample t-test). Values are represented as mean ± SEM, * p < 0.05.

**Supplementary Figure 7**

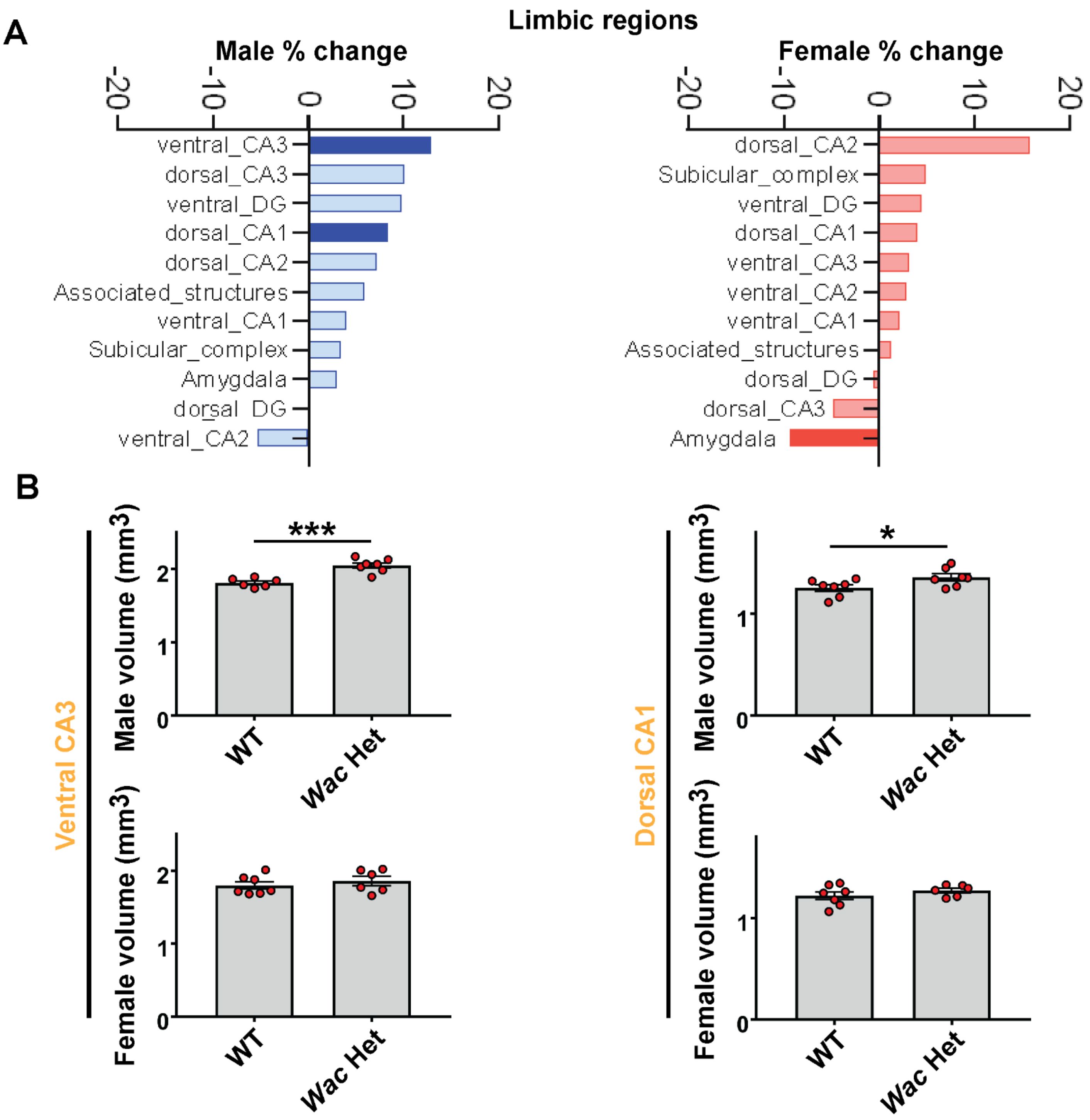


**Volumetric limbic regional changes**. (A) Percent change in limbic regions were modestly larger in HET mice than WT. In males the ventral CA3 and dorsal CA1 were larger but not in females. Female HET mice had smaller amygdala than WT female mice. (Bolded bars indicate significant differences p<0.05, t-test). (B) Ventral CA3 was significantly larger in HET male mice but not in female mice. Similarly, the dorsal CA1 also exhibited increased volumes in males HET mice but not WT nor in females. Data are expressed as the mean ± SEM; all groups n = 6-7, (*p < 0.05; ***p < 0.001).

**Supplementary Figure 8**

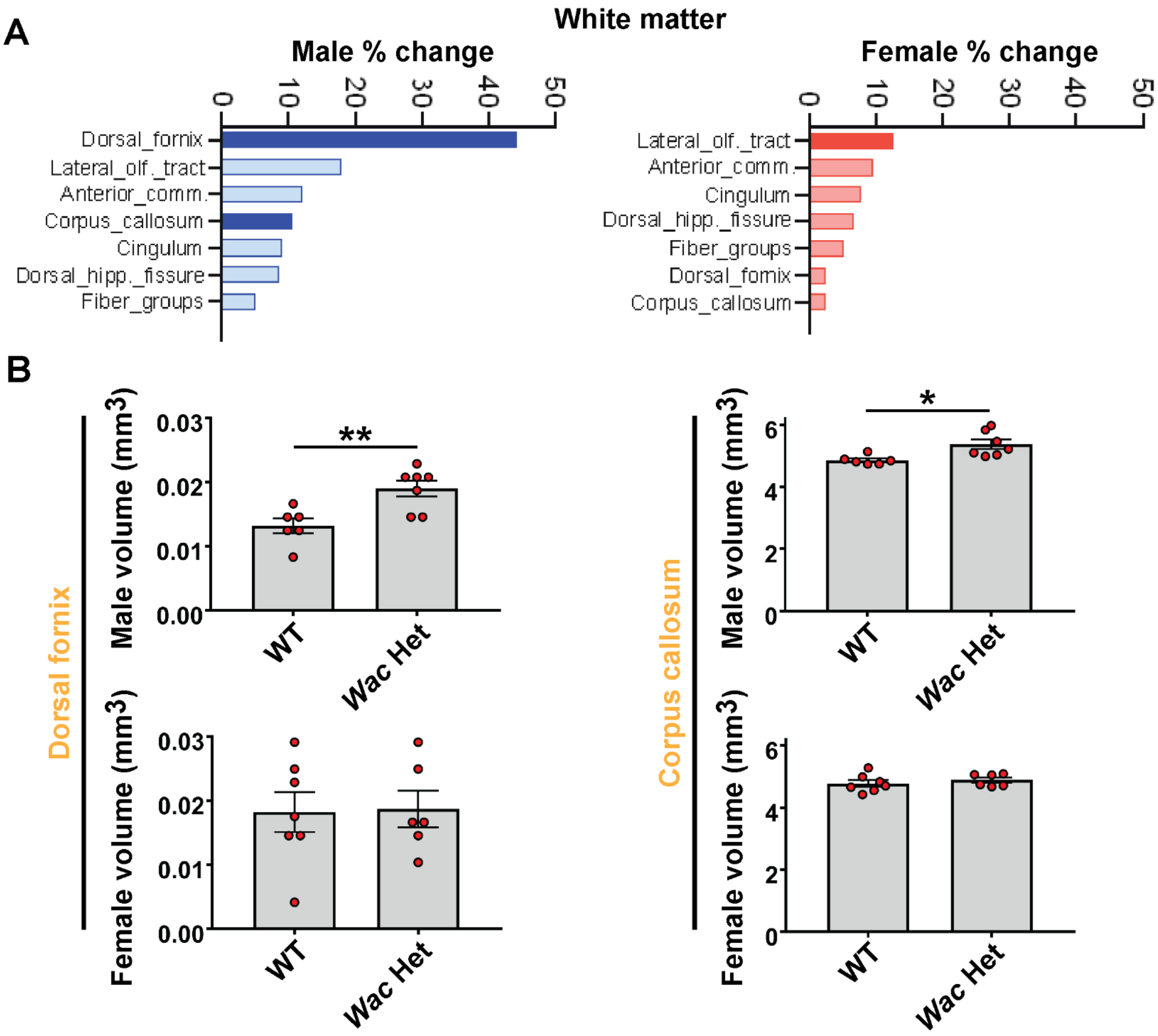


**Volumetric changes in white matter.** (A) Percent change in white matter regions were larger in male HET mice compared to WT, particularly the dorsal fornix. In females only the lateral olfactory tract was larger in HET mice than WT. (Bolded bars indicate significant differences p<0.05, t-test). (B) There was an increase in dorsal fornix volume in male but not female HET mice compared to WT. There was a robust increase in corpus callosum volume in HET male mice compared to WT but not in female HET mice. Data are expressed as the mean ± SEM; all groups n = 6-7, (*p < 0.05; **p < 0.01).

**Supplementary Figure 9**

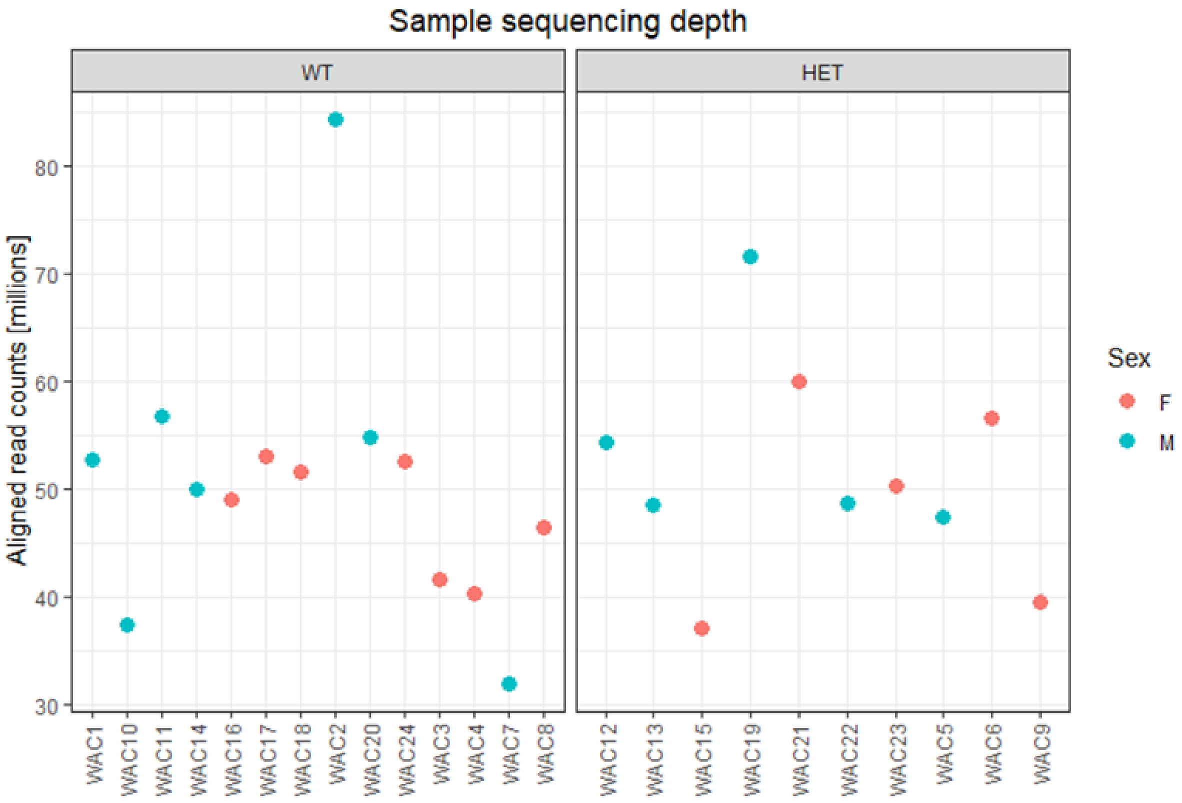


**Sample sequencing depth.** Values represent millions of aligned reads.

**Supplementary Figure 10**

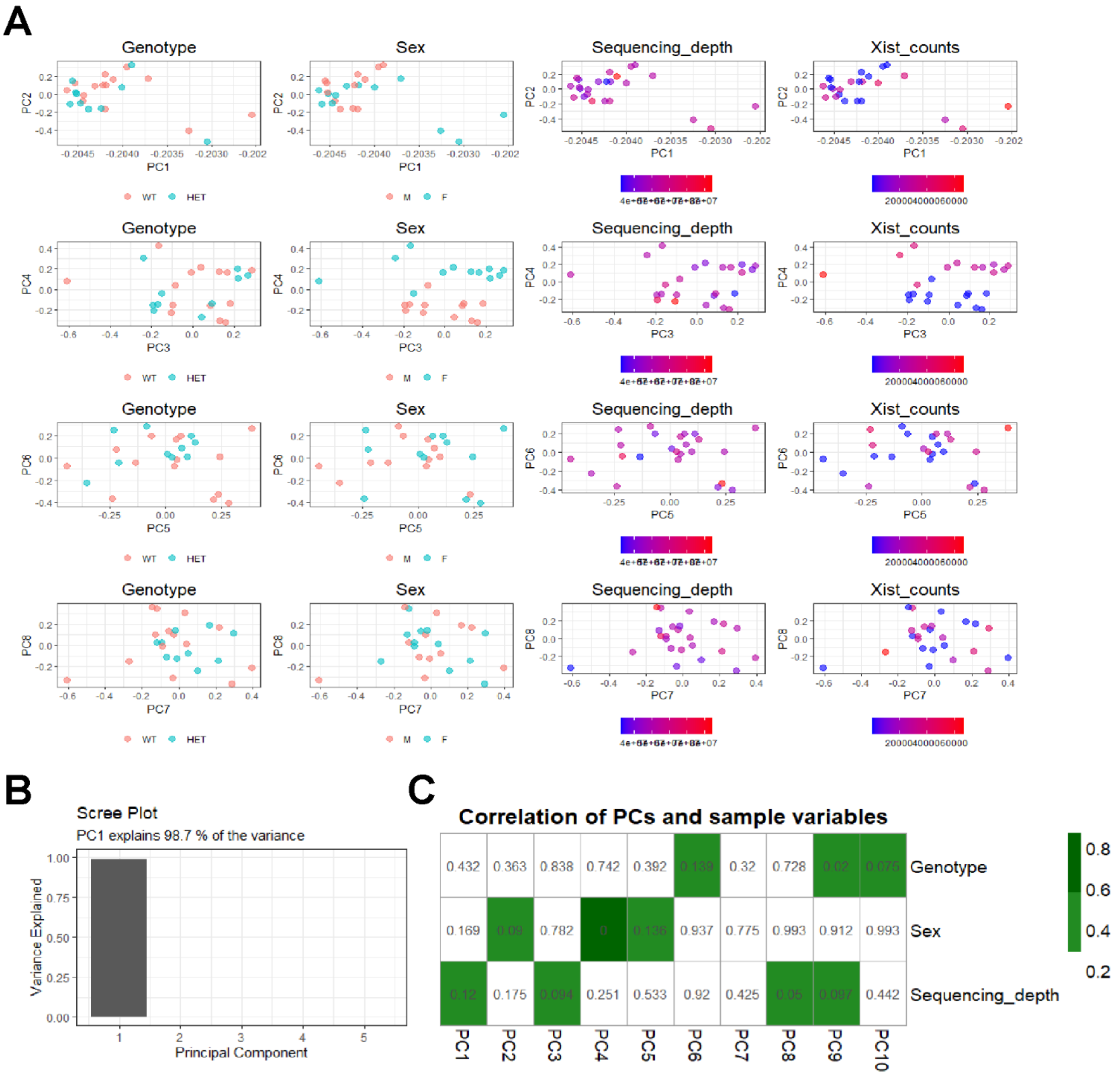


**Principal component analysis (PCA) dimensionality reduction.** (A) PCA sample clustering showing first 8 PCs. Samples are colored by genotype, sex, sequencing depth, and Xist gene counts. PCA did not identify sample outliers. (B) Scree plot showing the amount of variance explained by the first 5 PCs. (C) Correlation heatmaps of PCs 1 – 10 vs genotype, sex, and sequencing depth. Values in the heatmap represent correlation P values. The intensity of the heatmap colors represent Pearson correlation coefficient (r).

**Supplementary Figure 11**

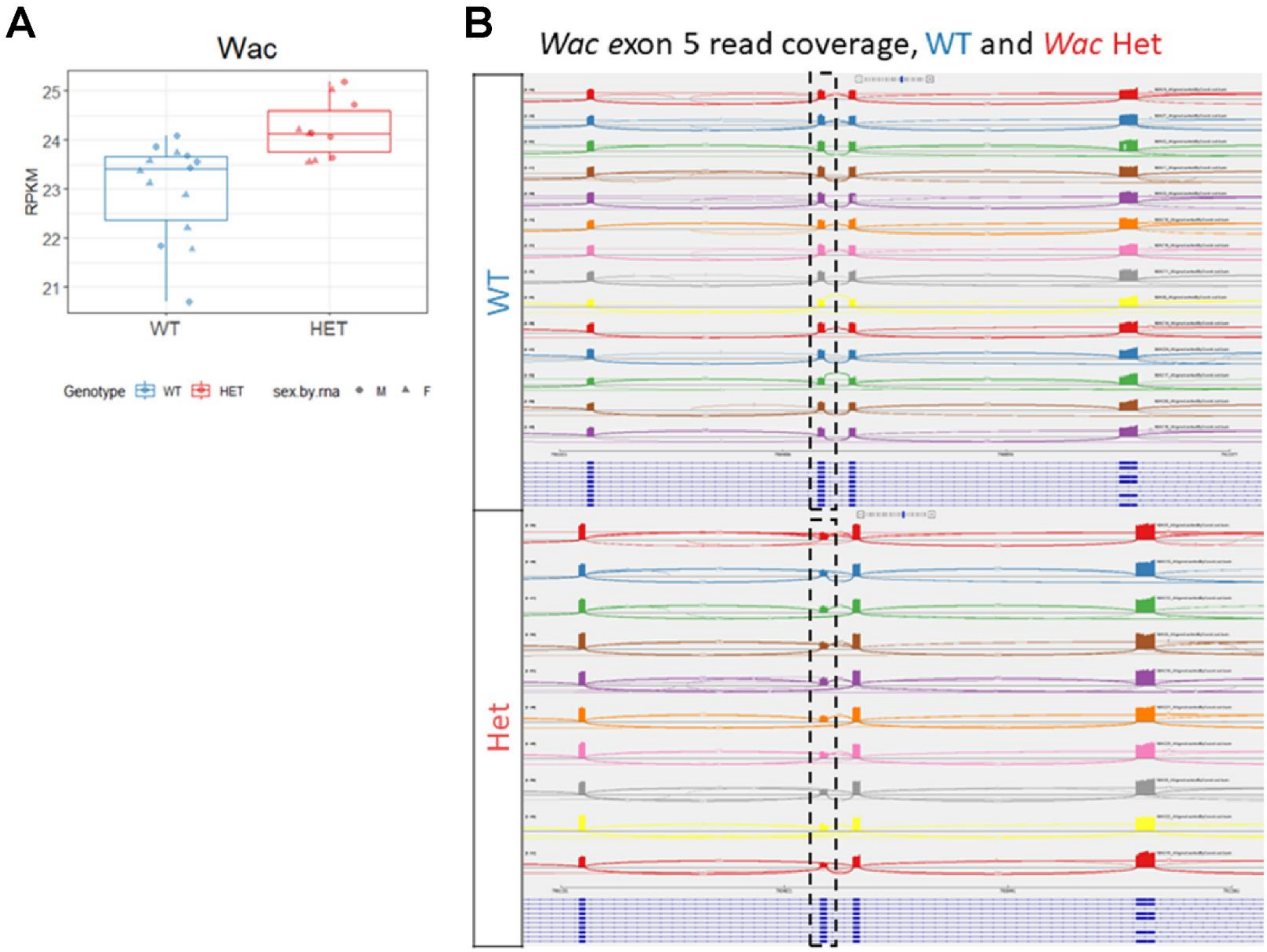


**Validation of *Wac* heterozygosity in RNA-seq data.** (A) *Wac* Het samples demonstrate subtle upregulation of *Wac* transcript, when reads are counted across the entire gene length (log2FC = 0.09, P = 0.01, FDR = 0.5) suggesting compensatory upregulation of *Wac* transcription. (B) Sashimi plots representing *Wac* read coverage. Dashed rectangle marks floxed exon 5, which coverage is reduced by ~50% in Het samples when compared to surrounding exons.

**Supplementary Figure 12**

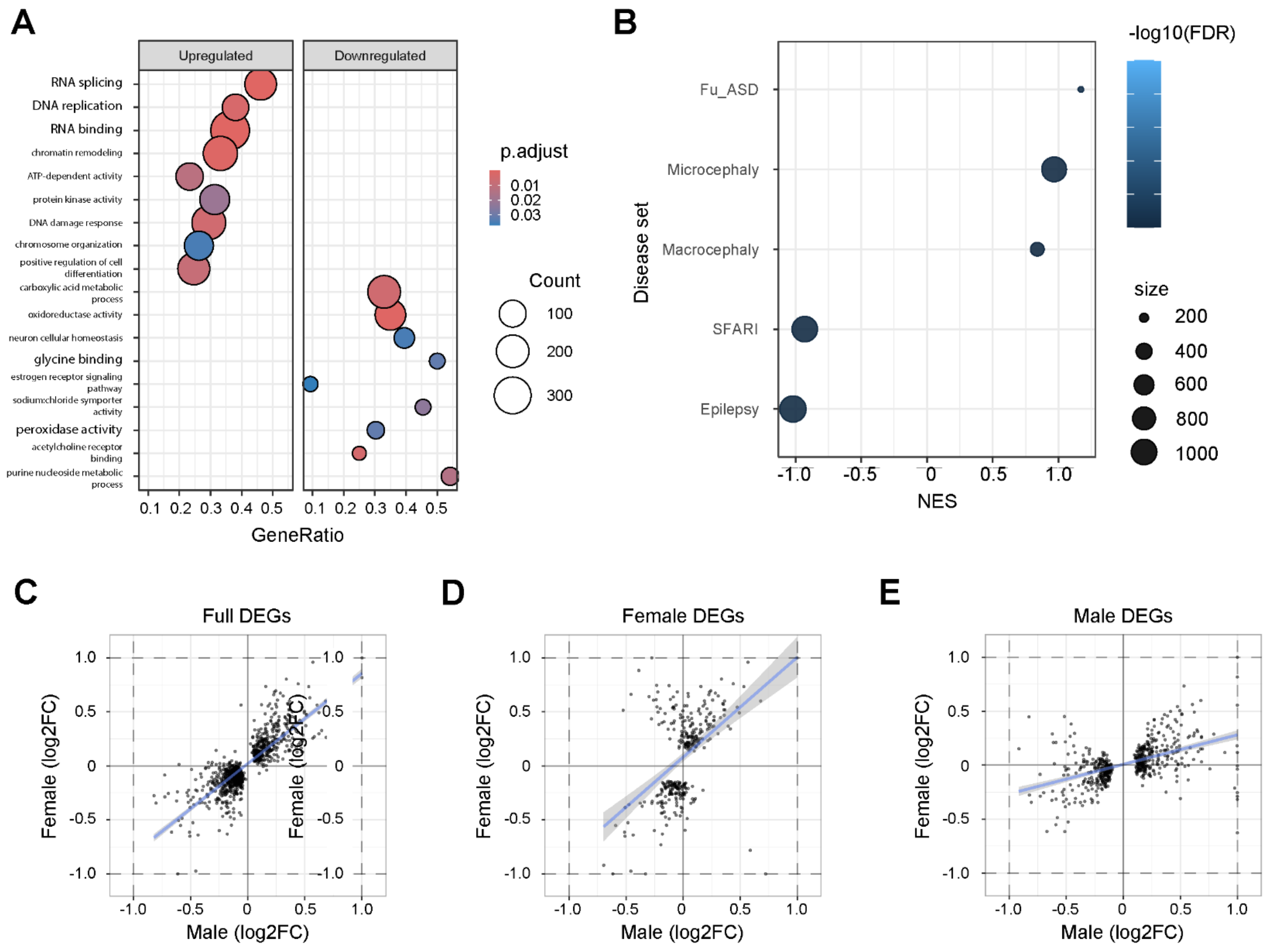


**Gene set enrichment and sex concordance in P2 *Wac* mouse forebrain bulk RNA-seq.** (A) Curated GO terms from SVA-corrected GSEA analysis. Colors indicate significance, size, the number of genes in the enriched term, and the enrichment score (NES). (B) Custom GSEA test looking for enrichment of disease associated genes in the SVA-corrected DE model. Colors indicate significance, size the number of genes in the enriched term, and the enrichment score (NES). (C) Correlation of the effected sizes (log fold change) of DEGs (pvalue < 0.05) identified in the SVA-corrected DE analysis between males and females. Line indicated a best fit linear regression with a 95% confidence interval. (D) Correlation of the effected sizes (log fold change) of DE genes (pvalue < 0.05) identified in the female DE analysis between males and females. Line indicated a best fit linear regression with a 95% confidence interval. (E) Correlation of the effected sizes (log fold change) of DE genes (pvalue < 0.05) identified in the male DE analysis between males and females. Line indicated a best fit linear regression with a 95% confidence interval.

### Supplementary Figure 13

A

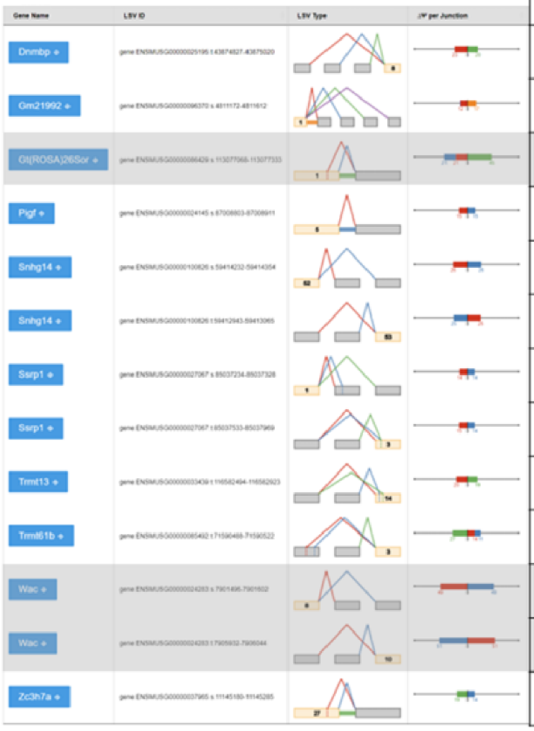


B

| Gene | Skipped exons (SE) | Alternative 5' (donor) splice sites (A5SS) | Alternative 3' (acceptor) splice sites (A3SS) | Retained introns (RI) | Mutually exclusive exon usage (MXE) |
|---|---|---|---|---|---|
| Dnmbp | | + | | | |
| Gm21992 | | | | + | |
| Gt(ROSA)26Sor | | | | + | |
| Pigf | | | | + | |
| Snhg14 | + | | | | |
| Snhg14 | + | | | | |
| Ssrp1 | | | + | | |
| Ssrp1 | | | + | | |
| Trmt13 | | | + | | |
| Trmt61b | | + | | | |
| Wac | + | | | | |
| Wac | + | | | | |
| Zc3h7a | | | | + (reduced in Het) | |

**Summary of differential splicing (DS) analysis.** (A) Visualizations of DS events, including splice graphs. Values in the ΔΨ per junction column represent percentage of the expected differential inclusion for matching edges in the splice graph. (B) DS event classification. DS events indicated in grey result from transgenesis.

### Supplementary Figure 14

WAC

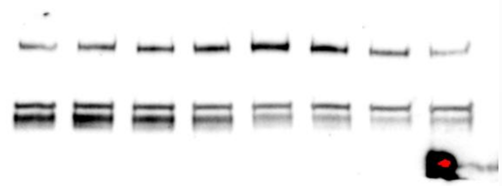

pS6

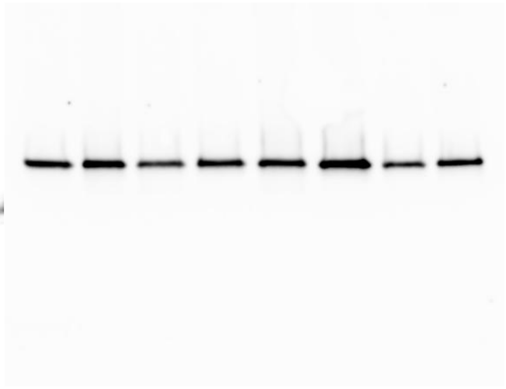

Total S6

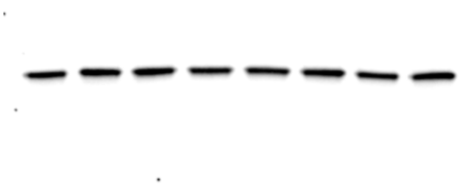

mUb-H2B

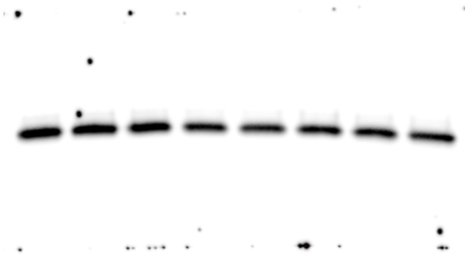

Beta-Catenin

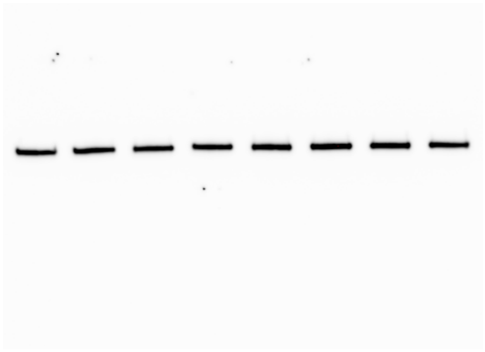

pERK1/2

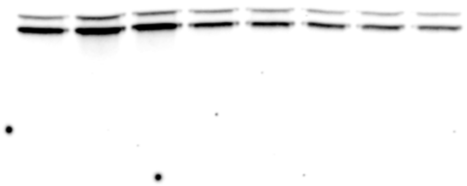

**Total ERK1/2**

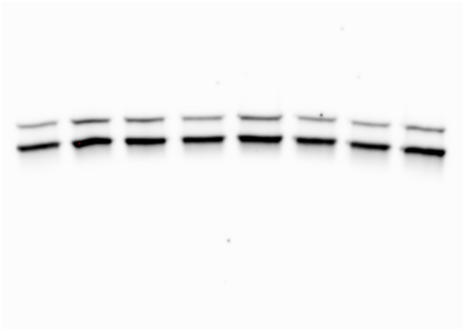

**PSD95**

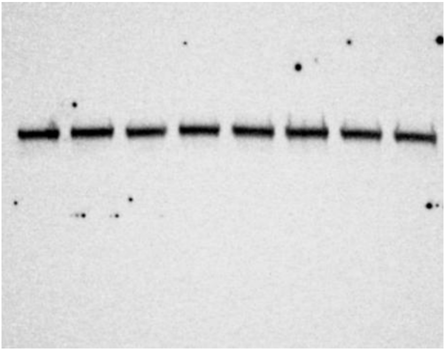

**GAD65/67**

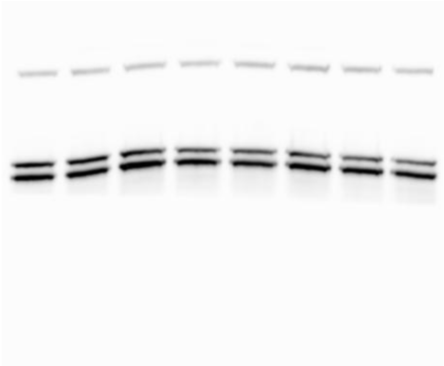

**Gephyrin**

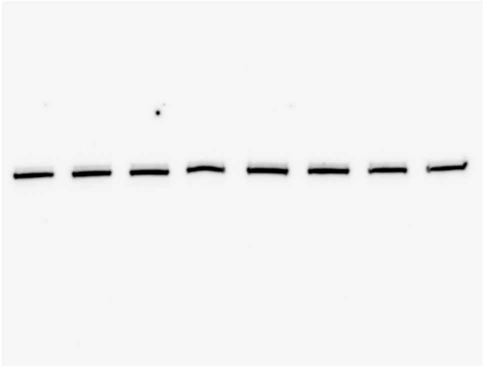

**GAPDH**

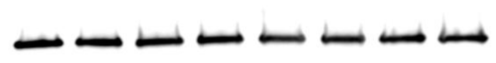

**alpha-Tubulin**

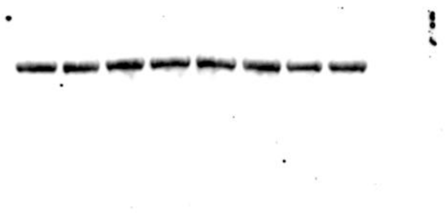

**Pan protocadherin alpha**

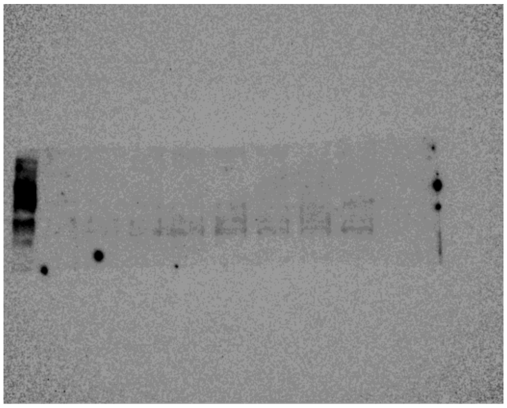

**Uncropped western blots.**

### Supplementary Table 1

| | *L2/3 excitatory neurons* | | | | | | | |
|---|---|---|---|---|---|---|---|---|
| | WT | | | *Wac* Het | | | | |
| | Mean ± s.e.m. | | | Mean ± s.e.m. | | | *P* | Test |
| RMP (mV) | -86.8 | ± | 0.7 | -87.1 | ± | 0.6 | 0.747 | *t*-test |
| $R_m$ (MΩ) | 66.8 | ± | 3.6 | 59.2 | ± | 3.7 | 0.151 | *t*-test |
| $\tau_m$ (ms) | 12.9 | ± | 0.5 | 12.3 | ± | 0.4 | 0.209 | *t*-test |
| $C_m$ (pF) | 197.4 | ± | 8.8 | 218.3 | ± | 13.7 | 0.720 | MW *U*-test |
| Rheobase (pA) | 170.0 | ± | 11.0 | 167.2 | ± | 12.6 | 0.868 | *t*-test |
| Spike threshold (mV) | -49.9 | ± | 0.6 | -50.1 | ± | 0.5 | 0.788 | *t*-test |
| Spike amplitude (mV) | 85.0 | ± | 1.5 | 88.1 | ± | 1.4 | 0.142 | *t*-test |
| Spike half-width (mV) | 0.63 | ± | 0.02 | 0.61 | ± | 0.02 | 0.323 | *t*-test |
| AHP amplitude (mV) | -18.3 | ± | 0.6 | -18.5 | ± | 0.5 | 0.794 | *t*-test |
| Max rate of rise (mV*ms) | 526.0 | ± | 28.8 | 586.3 | ± | 22.4 | 0.109 | *t*-test |
| Max rate of decay (mV*ms) | -112.8 | ± | 4.2 | -120.0 | ± | 4.5 | 0.243 | *t*-test |
| | n = 16 cells, 3 mice | | | n = 16 cells, 3 mice | | | | |

* See methods for an explanation of how the electrophysiological parameters were defined/measured.

* All membrane potentials were corrected for a 14 mV liquid junction potential

* All statistical tests were two-tailed

**Electrophysiological properties of L2/3 excitatory cells in the somatosensory cortex of WT and *Wac* Het mice**

### Supplementary Table 2

| ID | Genotype | Sex | RNA_concentration_ng_per_ul | RIN_score | RNA_230_260_ratio | Aligned_reads | Title |
|---|---|---|---|---|---|---|---|
| WAC1 | WT | M | 594.6 | 6.9 | 2.07 | 52762796 | WAC1_WT_M |
| WAC2 | WT | M | 765.3 | 9.2 | 2.09 | 84332816 | WAC2_WT_M |
| WAC3 | WT | F | 671.2 | 9.4 | 2.06 | 41664073 | WAC3_WT_F |
| WAC4 | WT | F | 588 | 9.3 | 2.03 | 40325503 | WAC4_WT_F |
| WAC5 | HET | M | 674.2 | 9.3 | 2.04 | 47396169 | WAC5_HET_M |
| WAC6 | HET | F | 599.3 | 9.3 | 2 | 56634354 | WAC6_HET_F |
| WAC7 | WT | M | 829.6 | 9.5 | 2.03 | 31992625 | WAC7_WT_M |
| WAC8 | WT | F | 712.1 | 9.5 | 2.02 | 46452921 | WAC8_WT_F |
| WAC9 | HET | F | 695.5 | 9.5 | 2.01 | 39539617 | WAC9_HET_F |
| WAC10 | WT | M | 926.8 | 9.5 | 2.03 | 37432155 | WAC10_WT_M |
| WAC11 | WT | M | 625 | 9.6 | 2 | 56705480 | WAC11_WT_M |
| WAC12 | HET | M | 527.5 | 9.5 | 2.2 | 54319711 | WAC12_HET_M |
| WAC13 | HET | M | 734.2 | 9.6 | 2.1 | 48516396 | WAC13_HET_M |
| WAC14 | WT | M | 724.8 | 9.5 | 2.07 | 50067506 | WAC14_WT_M |
| WAC15 | HET | F | 321 | 9.7 | 2.03 | 37146965 | WAC15_HET_F |
| WAC16 | WT | F | 621 | 9.6 | 2.04 | 49052221 | WAC16_WT_F |
| WAC17 | WT | F | 522 | 9.6 | 2.05 | 53045920 | WAC17_WT_F |
| WAC18 | WT | F | 579.8 | 9.8 | 2.04 | 51579332 | WAC18_WT_F |
| WAC19 | HET | M | 1062.7 | 9.7 | 2.06 | 71662005 | WAC19_HET_M |
| WAC20 | WT | M | 639.1 | 9.7 | 2 | 54832397 | WAC20_WT_M |
| WAC21 | HET | F | 970.4 | 9.6 | 2.09 | 59953779 | WAC21_HET_F |
| WAC22 | HET | M | 797 | 9.7 | 2.07 | 48767770 | WAC22_HET_M |
| WAC23 | HET | F | 1160.9 | 9.6 | 2.06 | 50341199 | WAC23_HET_F |
| WAC24 | WT | F | 916.6 | 9.7 | 2.07 | 52628428 | WAC24_WT_F |

**General parameters of the bulk-seq brains analyzed.** Columns indicate sample ID (ID), Genotye (WT = wildtype, HET = Wac mutant), Sex (M=Male, F= Female), RNA concentration, RNA integrity number (RIN) from bioanalyzer, A230/A260 value from spectrometer, number of aligned reads, and full sample metadata identifier (ID + Genotype + Sex)

**Supplementary Table 3:** SVA-corrected differential expression analysis of *Wac* mutants; combined sexes. Genes are annotated by common name and ENSEBLE gene id. Effect size indicated by log fold change (logFC) and relative expression by log counts per million (logCPM). Significance is reported by a likelihood ratio test statistic (LR), standard p-value (PValue), and the multiple-testing corrected p-value or false discovery rate (FDR).

**Supplementary Table 4:** Differential expression analysis of *Wac* mutants, using only male samples. Genes are annotated by both common name and ENSEBLE gene id. Effect size is indicated by log fold change (logFC) and relative expression by log counts per million (logCPM). Significance is reported by a likelihood ratio test statistic (LR), standard p-value (PValue), and the multiple-testing corrected p-value or false discovery rate (FDR).

**Supplementary Table 5:** Differential expression analysis of *Wac* mutants, using only female samples. Genes are annotated by both common name and ENSEBLE gene id. Effect size is indicated by log fold change (logFC) and relative expression by log counts per million (logCPM). Significance is reported by a likelihood ratio test statistic (LR), standard p-value (PValue), and the multiple-testing corrected p-value or false discovery rate (FDR).

**Supplementary Table 6:** GO over representation analysis of differential expression results for different models (Full, Male, Female). Ontology indicates the source of the term set (MF = Molecular Function, BP = Biological Processes, CC = Cellular Component). GO terms are identified by GO:ID number and term description. GeneRatio is the ratio of differentially expressed genes associated with the term to the total number of differentially expressed genes tested. BgRatio is the ratio of background genes associated with the term to the total number of genes in the background set. RichFactor is the ratio of observed differentially expressed genes to expected genes for the term (GeneRatio ÷ BgRatio). Fold enrichment of differentially expressed genes for the term relative to background expectation. The zscore, or standardized enrichment score reflecting deviation from expected overlap. Significance is indicated by the p-value from a Fisher's exact test, Multiple-testing–corrected p-value (p.adjust) using a Benjamini–Hochberg false discovery rate correction, and false discovery rate (qvalue). geneID shows the differentially expressed genes in the GO term. DE indicates direction of differential expression in the model. Model indicates the differential expression model (Full = SVA-corrected full dataset, Male = male only differential expression, Female = female only differential expression).

**Supplementary Table 7:** GO enrichment analysis using fgsea of differential expression results for different models (Full, Male, Female). Ontology indicates the source of the term set (MF = Molecular Function, BP = Biological Processes, CC = Cellular Component). GO terms are identified by GO:ID number and term description. SetSize indicates the number of genes from the gene set that are present in the ranked gene list used for GSEA. EnrichmentScore is the raw gene set enrichment score reflecting the degree to which genes in the set are overrepresented at the top or bottom of the ranked list. The normalized enrichment score (NES), which accounts for gene set size and allows comparison across gene sets. Significance is indicated by the p-value from a Fisher's exact test, Multiple-testing–corrected p-value (p.adjust) using a Benjamini–Hochberg false discovery rate correction, and false discovery rate (qvalue). Rank position at which the maximum enrichment score is observed in the ranked gene list. The leading edge is the subset of genes from the gene set that contribute most strongly to the enrichment signal. Core enrichment, an indicator or list specifying genes belonging to the leading-edge subset. Model indicates the differential expression model (Full = SVA-corrected full dataset, Male = male only differential expression, Female = female only differential expression).

**Supplementary Table 8: MAJIQ-based alternative splicing analysis of *Wac* mutants.** Local splicing variations (LSVs) are reported per gene and junction. Genes are annotated by Ensembl gene ID. Splicing effect size is quantified as the mean change in percent spliced in (mean ΔPSI) per LSV junction. Statistical confidence is reported as the posterior probability that the splicing event is changing (probability_changing) or non-changing (probability_non_changing). Mean PSI values are provided separately for wild-type (WT_mean_psi) and heterozygous mutants (Het_mean_psi). Additional structural annotations include the number of junctions and exons per LSV, genomic coordinates of junctions, and intron retention coordinates where applicable.

**Due to excessive length the above files are provided separately.** However, figure legends are provided next to each Table title above.

**Supplementary Table 9**

| Antibody (host/dilution) | Company | Category # | Clone | Lot # |
|---|---|---|---|---|
| Alpha-tubulin (mouse, 1:3000) | Abcam | ab7291 | DM1-A | GR310199-6 |
| Beta-catenin (rabbit, 1:4000) | Abcam | ab32572 | E247 | GR184212-83 |
| CTIP2 (rat, 1:500) | Abcam | Ab18465 | n/a | n/a |
| ERK1/2 (rabbit, 1:4000) | Cell Signaling | 4695 | 137F5 | 35 |
| pERK1/2$^{Thr202/Tyr204;Thr185/Tyr187}$ (rabbit, 1:4000) | Cell Signaling | 9101 | n/a | 31 |
| GAD65/67 (rabbit, 1:4000) | Sigma-Aldrich | G5163 | n/a | n/a |
| GAPDH (rabbit, 1:4000) | Cell Signaling | 2118 | 14C10 | 14 |
| Gephyrin (rabbit, 1:4000) | Cell Signaling | 43928 | D4J4R | 1 |
| IBA1 (goat, 1:500) | Cell Signaling | 17198 | E404W | 1 |
| LHX6 (mouse, 1:200) | Santa Cruz Bio. | sc-271433 | A-9 | B2219 |
| NeuN (mouse, 1:400) | Millipore | MAB377 | A60 | 3061189 |
| OLIG2 (rabbit, 1:400) | Millipore | AB9610 | n/a | 3071572 |
| Parvalbumin (rabbit, 1:400) | Swant | PV27 | n/a | 2014 |
| Protocadherin alpha (rabbit, 1:3000) | SySy | 190 003 | n/a | 1-8 |
| PROX1 (rabbit, 1:500) | Abcam | ab199359 | EPR19273 | GR3216583 |
| PSD95 (rabbit, 1:4000) | Cell Signaling | 3450 | D27E11 | 4 |
| S100beta (rabbit, 1:500) | Proteintech | 15146-1-AP | n/a | 00092556 |
| S6 (rabbit, 1:4000) | Cell Signaling | 2217 | 5G10 | 7 |
| pS6$^{Ser240/244}$ (rabbit, 1:4000) | Cell Signaling | 5364 | D68F8 | 8 |
| SATB2 (mouse, 1:500) | Abcam | Ab51502 | n/a | n/a |
| Somatostatin (rat, 1:200) | Millipore | MAB354 | YC7 | 2219310A |
| TBR1 (rabbit, 1:500) | Abcam | Ab31940 | n/a | n/a |
| mUb-H2B (rabbit, 1:4000) | Cell Signaling | 5546 | D11 | 6 |
| WAC (rabbit, 1:4000) | Abcam | ab109486 | n/a | GR175251-1 |

**Commercial antibodies and parameters used in this study.**
